# Ion-Pair–Free Nanoflow HILIC–MS With RNase Benchmarking for Native RNA

**DOI:** 10.1101/2025.11.18.689004

**Authors:** Yuyang Qi, Chengkang Li, Nur Yesiltac-Tosun, Jannick Schicktanz, Leona Rusling, Steffen Kaiser, Sam Wein, Stefanie Kaiser

**Author notes:** contributed equally to the manuscript (Yuyang Qi, Chengkang Li, Nur Yesiltac-Tosun).

## Abstract

RNA modifications play crucial roles in regulating cellular processes, but comprehensive mapping of the human RNome still remains limited by technological challenges. Mass spectrometry (MS) is a valuable tool to analyse RNA modifications complementing sequencing-based analysis. Current MS-based oligonucleotide workflows have limited sensitivity, requiring micrograms of RNA inputs and thus hindering studies on native RNAs. Additionally, environmentally toxic ion-pairing reagents are often required. Here, we report a highly sensitive, broadly applicable oligonucleotide-MS workflow that enables analysis of nanogram-scale RNA hydrolysates and we benchmark the substrate specificity of three nucleases: RNase T1, RNase 4, and colicin E5. We developed a nano-flow hydrophilic interaction liquid chromatography (HILIC) setup compatible with common MS buffers and coupled this with high-resolution MS. Using modified NucleicAcidSearchEngine (NASE), we confidently assigned RNA hydrolysates with diverse 3’-end chemistries. Furthermore, we demonstrate that RNase 4 and colicin E5 efficiently cleave modified RNAs including pseudouridine-containing transcripts, enabling high sequence coverages. Using this workflow, we successfully mapped modifications in 25 ng of native yeast tRNA^Phe^ and verified the sequence of 250 ng of a synthetic mRNA. Overall, our method provides a sensitive, high-resolution platform for oligonucleotide mass spectrometry, facilitating comprehensive analysis of RNA modifications and advancing efforts toward complete epitranscriptomic mapping.

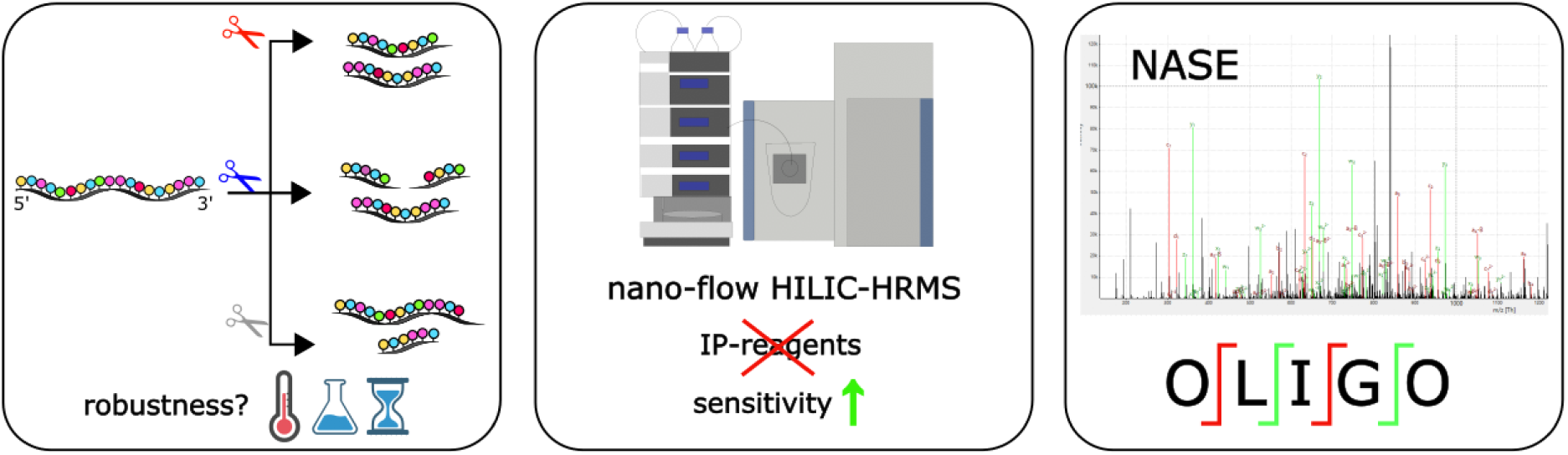

## Introduction

The central dogma of molecular biology states that DNA stores the genetic code, which is first transcribed into messenger RNA (mRNA) and then translated into proteins with the help of transfer RNA (tRNA) and ribosomal RNA (rRNA). Most of the RNAs investigated to date have been found to be decorated with chemical groups by special enzymes in a sequence-specific manner. These RNA modifications are summarized by the term epitranscriptome. To date, approximately 170 different RNA modifications are known (1), which occur in all life forms and RNA species studied to date. Depending on the RNA and modification, RNA modifications have different functions, which can be summarized by the term “life process regulation.” RNA modifications are recognized to impact human health (2), disease prevention (3), and agricultural success (4).

To exploit the full potential of RNA modifications and prevent or treat human diseases more efficiently, a deeper understanding of RNA modifications is needed. For this, a map of the “human RNome” is required, which, in addition to the sequence information of all cellular RNAs, also contains the chemical identity of the modification within its sequence context. This mapping urgently requires technological advances in RNA modification analysis.

The detection, localization, and quantification of RNA modifications are crucial for understanding the elusive function of these modifications. Two orthogonal concepts for RNA modification analysis have been established, namely mass spectrometry of RNA (RNA MS) and sequencing. Both have advantages and pitfalls which are partly complemented by the other technology. Especially short and highly modified RNAs remain a challenge for analysis. Next-generation sequencing (NGS) relies on reverse transcription (RT) of RNA into cDNA followed by amplification, allowing sensitive detection of nanogram amounts of RNA. However, most RNA modifications are lost during cDNA synthesis, and only those that block RT can be detected (5). Third-generation sequencing, such as Oxford Nanopore Technology, sequences RNA directly by measuring changes in electrical current as nucleotides pass through a pore. While it avoids cDNA synthesis and has the potential to detect modifications directly, RNAs should be at least 50 nts long for analysis and extensive validation is required (6,7).

Mass spectrometry is a valuable tool to analyse RNA modifications, and it is the only technology that allows chemical characterization of RNA modifications (8). Commonly, RNA is purified and hydrolysed to the nucleoside building block with subsequent targeted MS analysis. While this approach provides valuable information about the absolute abundance and stoichiometry of RNA modifications, its key bottleneck is the loss of all sequence information.

Bottom-up (or shotgun) oligonucleotide MS, the MS analysis of partial hydrolysates, is a reasonable method to overcome the loss of sequence information while retaining the chemical identification of a modification. First reports of bottom-up RNA MS date back into the 90’s when electrospray ionisation (ESI) entered the mass spectrometry field (9). The bottom-up approach relies on four key steps, namely RNA hydrolysis into short RNA fragments (5-15 nts), chromatographic separation of the resulting oligonucleotides, mass spectrometric detection, and data analysis. Nowadays, oligonucleotide MS allows complete mapping of RNA modifications onto synthetic mRNA sequences (10), native tRNA mixtures, and native rRNAs (11–14).

The bottleneck of oligonucleotide MS is its low sensitivity and thus the need of micrograms of purified RNA is the key limitation. Additional limitations of the technology are (a) ion-pairing reagents for oligonucleotide separation that are needed due to high hydrophilicity of oligonucleotides (IP-RP-LC), (b) the need for RNases with higher specificity to create longer fragments that are unique (e.g. 15-30 nts), (c) isomers such as U and Ψ, m^1^A and m^6^A or m^1^G, m^2^G and m^7^G that cannot be easily distinguished. For these reasons, oligonucleotide MS is not regularly used on native RNA isolates from *e.g.* human or bacterial cells.

Here, we show our recent progress on four key aspects of oligonucleotide MS and we combine the optimized parameters for analysis of nanogram amounts of RNA hydrolysates. Namely, we systematically assessed the substrate specificity of the dinucleotide specific RNases colicin E5 and RNase 4 and benchmarked these against the known RNase T1. We determined the robustness of substrate specificity in dependence of enzyme:RNA ratio, incubation time, temperature, and buffer composition; and we compared specificity using MALDI-MS and oligonucleotides-MS. In a next step, we developed a HILIC nanoflow chromatography which utilizes ammonium acetate, water, and acetonitrile, making it compatible with commonly applied buffers in MS core facilities. In conjunction with a self-packed diol-chemistry stationary phase, we obtain excellent chromatographic resolution. Using a qExactivePlus mass spectrometer, we analyzed unmodified RNA hydrolyzed with either RNase 4, colicin E5, or RNase T1. For data analysis, we used nucleic acid search engine (NASE) (15) and BioPharmaFinder; we found NASE to be superior in analysis of small, densely modified RNAs. Finally, we combined all our efforts, and we succeeded to map RNA modifications in 50 ng of native tRNA^Phe^ from yeast and verify the sequence of a synthetic mRNA using only 250 ng. In conclusion, we now have a broadly applicable and sensitive workflow for analysis of native RNAs from cellular lysates.

## Materials & Methods

### Chemicals and Reagents

All chemicals and reagents were purchased from Sigma Aldrich (St. Louis, MO, USA) unless otherwise specified.

RNA and DNA oligonucleotides were purchased from Sigma-Aldrich or Dharmacon, as specified in Table S1.

Firefly luciferase (FLuc) mRNA was purchased from NIST (The National Institute of Standards and Technology, RGTM 10202).

LC-MS grade acetonitrile (ACN) (Art. No. HN40), Urea (3941.1) and Tris (4855.2) were obtained from Carl Roth (Karlsruhe, Germany). LC-MS grade ammonium acetate (84885.180P, HiPerSolv CHROMANORM®) and Acetic acid (84874.180, HiPerSolv CHROMANORM®) were purchased from VWR (Darmstadt, Germany).

### Expression and purification of colicin E5

The expression plasmid pD454-CE5-Im5 (GenScript Biotech, Netherlands) was transformed into BL21(DE3) chemically competent cells. Transformed cells were plated on LB-agar containing 50 µg/mL kanamycin and incubated overnight at 37°C. A single colony was inoculated into TB medium supplemented with chloramphenicol (0.034 mg/mL) and kanamycin (0.1 mg/mL) and incubated overnight at 37°C. For large-scale expression, the culture was expanded, and protein overexpression was induced with 0.5 mM IPTG at OD600 = 1.5, followed by overnight incubation at 18°C. Cells were harvested by centrifugation (6000 rpm, 4°C, 15 min) and resuspended in lysis buffer (25 mM HEPES, pH 7.5, 500 mM NaCl, 25 mM imidazole, 5% glycerol, 0.5 mM TCEP). After sonication, nucleic acids were removed by polyethyleneimine (PEI) precipitation, and the lysate was cleared by centrifugation (23,000 × g, 40 min). The supernatant was filtered (0.45 µm) and loaded onto a 5 mL IMAC column (ÄKTAprime system) pre-equilibrated with lysis buffer. The column was washed with 10 mM imidazole buffer, followed by sequential washes with refold buffer 1 (25 mM HEPES, pH 7.5, 25 mM NaCl, 5% glycerol, 0.5 mM TCEP) and refold buffer 2 (25 mM HEPES, pH 7.5, 500 mM NaCl, 5% glycerol, 0.5 mM TCEP). CE5 was eluted using a 0–300 mM imidazole gradient in IMAC elution buffer (25 mM HEPES, pH 7.5, 500 mM NaCl, 300 mM imidazole, 5% glycerol, 0.5 mM TCEP). For further purification, CE5-containing fractions were pooled, concentrated, and subjected to size-exclusion chromatography (SEC) using a Superdex™ 75 column in SEC buffer (20 mM HEPES, pH 7.5, 250 mM NaCl, 0.5 mM TCEP). The purified colicin E5 protein was concentrated to 3.5 mg/mL, yielding ∼20 mg, and stored at –80°C.

### In vitro transcription (IVT) *of E. coli* tRNA^Ile^

Plasmid *pPK1204* with the correctly sequenced tRNA^Ile^ insert was transformed into *E. coli* DH5α competent cells and cultured on a large scale to produce sufficient plasmid DNA. The plasmid DNA was isolated using a Qiafilter Plasmid Maxi Kit (Qiagen, Venlo, Netherlands) and linearized with *HindIII* according to manufacturer protocol. *In vitro* transcription of tRNA^Ile^ was conducted using optimized concentrations of DNA template and magnesium acetate. For large-scale transcription, the process was performed under these optimized conditions for a duration of 4 hours. Specifically, the conditions included 40 mM magnesium acetate and 100 ng/µL of DNA template. RNA products, including HDV ribozymes (Hepatitis-Delta-Ribozyme) and uncut RNA, were separated by preparative urea-polyacrylamide gel electrophoresis (PAGE). tRNA^Ile^ was then extracted from the gel, ethanol precipitated, and desalted using a PD10 column (Cytiva, Marlborough, Massachusetts). The RNA was concentrated using a SpeedVac (Thermo Fisher Scientific, Waltham, Massachusetts) and subsequently stored at - 20°C.

### RNase T1 digestion of tRNA into oligonucleotides

For RNase T1 digestion, RNase T1 (Thermo Fisher Scientific, 1000 U/µL) was diluted in 25 mM Tris-HCl (pH 7.5) and 100 mM NaCl. Up to 1 µg of RNA was digested with RNase T1 at 37°C for 30 minutes in a total reaction volume of 50 µL, maintaining final concentrations of 25 mM Tris-HCl (pH 7.5) and 100 mM NaCl. For the IVT and biological samples, a final ratio of 10 U RNase T1 per µg of RNA was used.

### RNase 4 digestion of tRNA into oligonucleotides

The protocol was adapted from Wolf et al. (16). RNA was initially denatured by adding 3 M urea (Carl Roth, Germany) to achieve a final concentration of 1 M. The sample was then incubated at 90°C for 10 minutes, followed by immediate transfer to and maintenance at 37°C. Subsequently, the cooled RNA solution was diluted threefold in NEBuffer r1.1 (New England Biolabs) by adding twice the volume of 1.5× NEBuffer r1.1, which had been pre-warmed to 37°C. Human RNase 4 (New England Biolabs) was pre-diluted to the desired concentration, and 1 µL of the diluted enzyme was added to the RNA mixture. The reaction was incubated at 37°C for 1 hour. For the IVT and biological samples, a final ratio of 20 U RNase 4 per µg of RNA was used.

### Colicin E5 digestion of tRNA into oligonucleotides

For colicin E5 digestion, 1 µg of in-vitro transcribed tRNA was incubated with the C-terminal domain of colicin E5 at a molar ratio of 1:4 (RNA:colicin E5) in a reaction buffer containing 25 mM Tris-HCl (pH 7.5) and 100 mM NaCl. The total reaction volume was adjusted to 50 µL with nuclease-free water. The reaction mixture was gently mixed and incubated at 37°C for 30 minutes.

### Polyacrylamide Gel Electrophoresis

Oligonucleotides were analysed using 20% TBE-urea PAGE. Before sample loading, the gel was pre-run at 250 V for 30 min for equilibration. Samples (20 pmol) were mixed 1:1 with 2× loading buffer and loaded into wells (20 µL per sample). A molecular ladder containing 10 pmol of oligonucleotides (5-, 8-, 10-, 20-, 30-, 40-mer, tRNA^Ile^, HDV) in 2× RNA loading dye (NEB, Ipswich, MA, USA) was prepared and loaded similarly. Electrophoresis was performed at 275 V for 60–90 min. The gel was stained for 10 min with StainsAll solution (0.65× TBE, 0.01% StainsAll, 10% formamide, 25% isopropanol, 65% water), destained for 2 h in 1× TBE buffer containing 25% isopropanol and imaged using a Bio-Rad ChemiDoc™ MP Imaging System.

### Sample preparation for MALDI-MS and oligonucleotide-MS

To prevent RNA over-digestion, samples were filtered immediately after RNase-digestion using Zymo-Spin filters (C1004-50, Zymo Research Europe GmbH, Freiburg, Germany) according to the manufacturer’s instructions. Following incubation, the reaction mixture (50 µL) was combined with 100 µL of Zymo oligo-binding buffer. Subsequently, 400 µL of absolute ethanol was added, and the resulting solution was mixed thoroughly. The entire mixture was loaded onto a Zymo-Spin column and centrifuged at 12,000 rcf for 30 seconds. After discarding the flow-through, 750 µL of DNA wash buffer was added to the column, followed by a second centrifugation at 12,000 rcf for 60 seconds. The Zymo-Spin column was then transferred to a clean microcentrifuge tube, and 6-10 µL of RNase-free water was carefully applied to the center of the column membrane. To elute the purified oligonucleotide fragments, the column was subjected to a final centrifugation at 12,000 rcf for 60 seconds. The eluted RNA was stored for further analyses by MS.

### MALDI-TOF mass spectrometry of oligonucleotides

For each sample, 0.5 µL of 3-HPA matrix (3-hydroxipicolinic, half-acid saturated in H_2_O:Acetonitrile (1:1; v:v) (Bruker Daltonics, Bremen, Germany), containing 10 mg/mL diammonium hydrogen citrate) were spotted on an AnchorChip (800 µm) target. The matrix solution was allowed to dry at room temperature. Then 0.5 µL of each sample (7.75 µM) were spotted on top of the dried matrix preparation spot. The sample was allowed to dry and the spectra were acquired in positive ion mode with reflector on an ultrafleXtreme MALDI-TOF-TOF mass spectrometer (Bruker Daltonics, Bremen, Germany).

### Liquid chromatography (tandem) mass spectrometry (LC-MS)-based oligonucleotide analysis

*Materials* used for LC development: Thermo RP column: Acclaim™ PepMap™ 100 C18 HPLC column (150 mm x 0.075 mm, 2 μm, 100 Å, cat# 164534), RP trap cartridge: PepMap™ Neo Trap Cartridge, (5 mm x 0.3 mm, 5 μm, 100 Å, cat#: 174500) and Thermo amide: Accucore™ 150 Amide HILIC HPLC column (150 mm x 0.075 mm, 2.6 μm, 150 Å, cat# 16726-157569, Thermo Scientific Inc., Karlsruhe, Germany). HILIC stem trap cartridge: EXP2 0.33 μL stem trap (13.5 mm x 180 μm, 5 μm HALO PELL HILIC, 90 Å, P/N: 15-04001-ES, Optimize Technologies, Oregon, USA). Self-pack diol column: 150 mm x 0.075 mm, 3 μm, 120 Å, P/N: r13.d0.0001, Dr. Maisch HPLC, Ammerbuch, Germany and empty capillary: 150 mm x 0.075 mm.

*Sample Preparation* was performed using Zymo spin columns as outlined in the section above. To 0.8 µL RNA, 3.2 µL of acetonitrile was added for all injections and 1 µL was injected. To showcase the impact of RNA hydrolysate injection with 100% aqueous vs. 20% aqueous, the addition of acetonitrile was omitted once.

The LC-MS-based oligonucleotide analysis was performed on a UHPLC Ultimate™3000 RSLCnano system (Thermo Fisher scientific, Germering, Germany) coupled with a Q Exactive Plus hybrid quadrupole-Orbitrap mass spectrometer. For the Q Exactive Plus measurements, a Pepsep sprayer (P/N PSS1, Bruker, Billerica, MA, USA) was paired with a Pepsep integrated liquid junction stainless steel emitter (P/N 1893525, 30 μm ID, Bruker, Billerica, MA, USA).

Samples were injected either via the “direct” or “pre-concentration” injection modes during method optimization using 80/20 (v/v) H_2_O/ACN with 15 mM ammonium acetate (pH 5.5) as mobile phase A (MPA) and 20/80 (v/v) H_2_O/ACN with 15 mM ammonium acetate (pH 5.5) as mobile phase B (MPB) (12). Samples were pre-diluted in 80% ACN before injection. While the “pre-concentration” injection mode was conducted, samples were firstly trapped onto a EXP2 0.33 μL stem trap using MPB for 5 min at a 5 μL/min flow rate, unless otherwise mentioned. Afterward, the pre-concentrated samples on trap were reverse-eluted onto the nano analytical column depending on the experiment using a 0.3 μL/min flow rate at 30 °C, unless otherwise stated. The LC gradients for different experiments are shown in the respective section. Blank injections (i.e. MilliQ water) were included between every sample run. UV absorption at 260 nm was recorded throughout the elution.

#### Settings on the Q Exactive Plus

Ionization was achieved at 1.8 kV spray voltage in the negative ion mode at 250 °C and the MS was set to “Full MS/data-dependent MS^2^ (dd-MS^2^)” method with a scan range of 500-2500 *m/z*. Resolution, AGC target, and maximum injection time for both MS and MS^2^ were set to be 70,000 (FWHM), 1e6, and 300 ms, respectively. The Top 15 most intense ions were selected for MS^2^ fragmentation in the higher-energy collisional dissociation cell (HCD) using the normalized collision energy (NCE) set to be 28 with 10 s dynamic exclusion period with *m/z* 1.7 isolation window width.

### NucleicAcidSearchEngine (NASE) analysis

The LC-MS raw data files were converted to mzML format files using MSConvert (https://github.com/ProteoWizard/) and analyzed using the NASE software (OpenMS ver. 3.0.0-pre-nightly-2023-03-09, https://openms.de/) for database matching (15,17,18).

Target/decoy database searches were performed with 5% false discover rate cutoff. Mass tolerances for precursor and fragment ions were set to be 10 ppm, respectively, while precursor charge states of -2 to -20 were included. Potential cation adducts, including sodium (Na^+^), potassium (K^+^), and ammonium (NH^4+^), were included in the search for their potential contribution to the shift of precursor *m/z* values. Precursor isotopes of -1, 0, 1, 2, and 3 were included, while all possible fragment ion types, including a-B, a, b, c, d, w, x, y, and z ions, were selected in the searches. The detailed configuration used for all our data analysis procedures is specified in Figure S19, where the corresponding nuclease must be chosen in the option “enzyme”. The NASE analysis of synthetic ribonucleotides and poly-dT oligonucleotides were manually verified in Skyline software (ver. 22.2.0.351, University of Washington, Seattle, WA, USA) using self-prepared transition list, including retention time, isotopic correlation, and precursor mass tolerance (19).

### Use of an AI language model

We used an AI assistant (GPT-5 Thinking) to support manuscript preparation in well-defined, author-directed tasks. Specifically, we prompted the model to: (i) extract the transcribed FLuc mRNA sequence from the provided PDF and generate a FASTA file and (ii) language editing for clarity, logical flow, structure, concision, terminology harmonization, and grammar. All AI-generated outputs were reviewed, edited, and verified by the authors; data analysis decisions, figures, and text content remain the responsibility of the authors.

## Results

### Comparison of RNase activity

To verify the activity of the purchased RNases T1 and 4 and our purified colicin E5, we first performed a cleavage assay using an *in-vitro* transcribed tRNA^Ile^ as a benchmark RNA for the following experiments. The sequence of tRNA^Ile^ contains several G’s, UG/UA and GU motifs, allowing us to follow potential cleavage events at different sites for RNase T1, RNase 4 and colicin E5 (Figure 1A). 1 µg of RNA was incubated with increasing amounts of each RNase according to the respective RNase’s standard protocol and the RNA was analysed on a 20% denaturing gel. We verified the activity of the enzymes, as the full-length tRNA band disappeared after treatment with the enzymes and we obtained product bands below the tRNA band (Figure 1B-D). As expected, with lower amounts of enzyme we received longer fragments, potentially due to missed cleavages. We then subjected all samples to MALDI-TOF MS analysis and determined the sequence of the resulting fragments. An exemplary spectrum of an intermediate enzyme concentration is found for each RNase in Figure 1B-D and for all other concentrations in Figures S1-3. The MALDI spectra change towards smaller *m/z* values with rising RNase T1 concentrations which is in accordance with expectation due to the decreasing number of missed cleavages. Furthermore, we observe a dependence of the type of 3’ end on the RNase concentration. Especially at lower enzyme concentrations, we observe the presence of 3’-cyclic phosphate intermediates, consistent with the two-step catalytic mechanism of RNase T1 involving a 2’,3’-cyclic phosphates intermediate and a slower hydrolysis step to a 3’-phosphate. These findings are consistent with previous studies (20) indicating that enzyme concentration and temperature can influence the ratio of cyclic versus linear 3’-phosphate products. At higher RNase T1 content, we observe primarily 3’-phosphate ends. For RNase 4, we also observe changes in the MS spectra in dependence of the studied RNase concentration. Unlike with RNase T1, we do not see smaller fragments with rising enzyme concentration but rather different fragments under the different conditions. This, and the fact that we could not assign all fragments according to the expected U|A and U|G cleavage site, indicates that the substrate specificity of RNase 4 is more variable. At higher enzyme-to-substrate ratios, we additionally detected an increased frequency of C|A cleavage sites, consistent with previous findings (21). This highlights the importance of applying moderate enzyme-to-substrate ratios when specific cleavage patterns are desired. Notably, RNase 4 predominantly generates 2’,3’-cyclic phosphate ends. On the gel we noticed a long fragment around 35 nts, which is probably the result of a missed cleavage in a double-stranded stretch of the tRNA substrate. This fragment could not be detected in the MALDI spectrum as we could not ionize fragments exceeding 25 nts. For colicin E5, all MALDI spectra were similar and did not depend on the chosen concentration of enzyme. Most signals in the MALDI spectrum could be assigned to the expected G|U cleavage site and most fragment signals were with a 2’,3’-cyclic phosphate group, rather than a 3’-phosphate.

**Figure 1.**
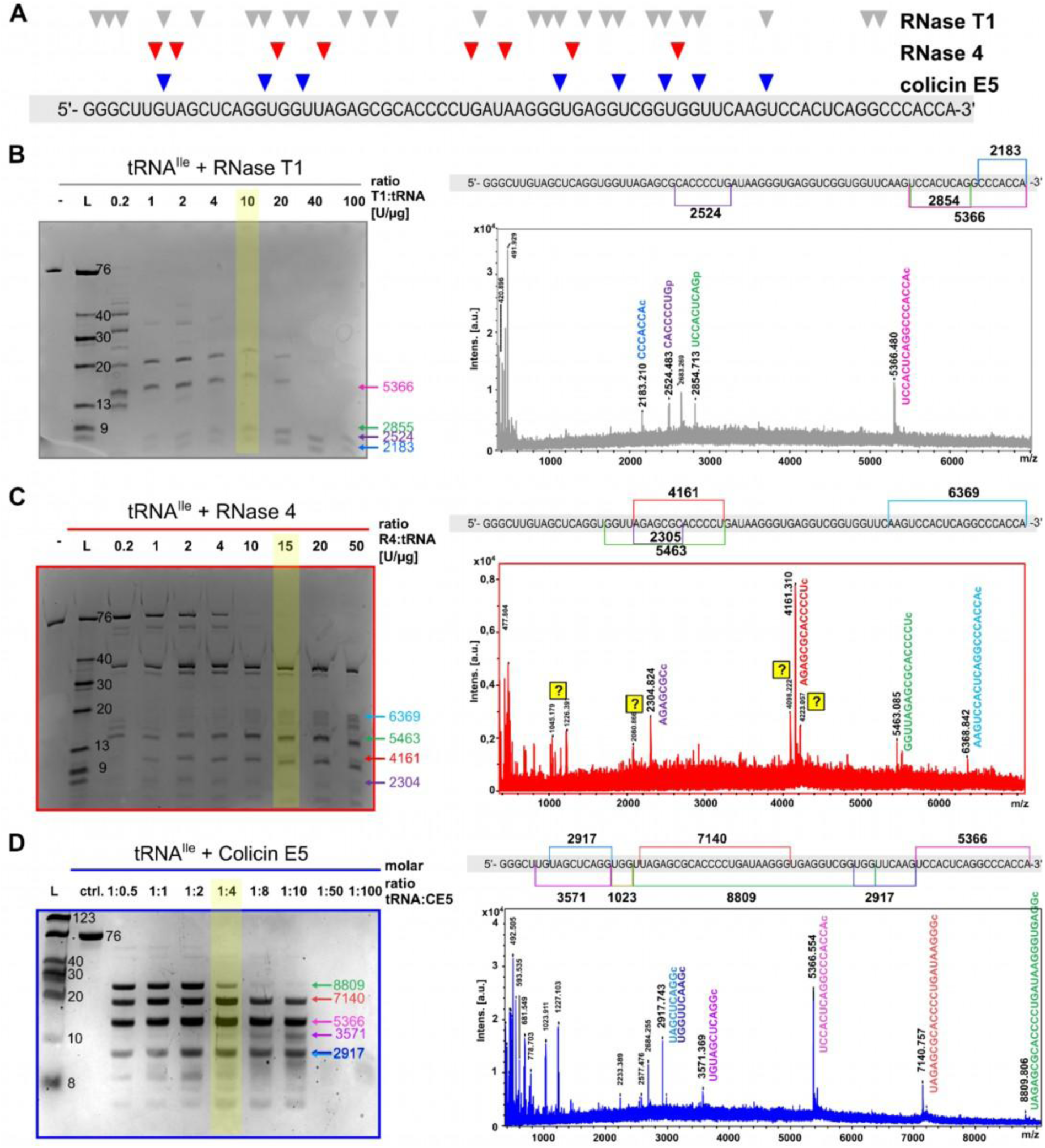
Substrate specificity of common nucleases for partial RNA hydrolysis. (**A**) sequence of an unmodified tRNA (*E. c.* tRNA^Ile^_GAU_) and the expected cleavage sites of three common RNases. RNase T1 – grey, RNase 4 – red, colicin E5 – blue. (**B**)-(**D**) polyacrylamide gel of *E. c.* tRNA^Ile^_GAU_ treated in different ratios with the three RNases and the corresponding MALDI-MS spectra. Assigned fragments and cleavage sites are added to the MS spectrum and in the sequence above. Digestion conditions: 37 °C, 30 minutes for RNase T1 and colicin E5 and 60 minutes for RNase 4. Arrows in the gel give the [M+H]^+^ of the indicated bands.

All in all, we find that all three enzymes can produce oligonucleotides of the length needed for further analysis of native RNA. Yet, we find a high variability either in the 3’-end or cleavage site specificity for all three RNases, which complicates downstream data analysis. To minimize this effect, we systematically assessed the influence of the reaction conditions on product variability.

### Variability of reaction products is only mildly influenced by the reaction conditions

To further assess the impact of the enzymatic digestion conditions on the resulting oligonucleotides’ chemistry, we defined “starting” parameters for each nuclease (Table S2) and varied the incubation time, temperature, buffer composition (including magnesium ions), and pH. Oligonucleotide length was assessed by gel electrophoresis. As shown in Figure S4, reaction products generated by colicin E5 remained highly similar across all tested conditions. For RNase T1 and RNase 4, temperature variation had only a moderate effect on product formation. Both enzymes were active even on ice, reaching maximum activity around 37 °C and 50 °C, where no fragments longer than 20 nt (RNase T1) and no intact tRNA (RNase 4) were detectable. At 50 °C, further shortening of the fragments was observed. Increasing the temperature to 75 °C, however, inhibited enzymatic activity, leading to the appearance of longer fragments. Yet, residual activity remained, confirming that heat inactivation at 75 °C is not effective.

Reaction kinetics were rapid for all three nucleases, with RNase T1 and RNase 4 producing distinct cleavage products within 5 minutes. Prolonged incubation led to the generation of shorter fragments.

We further compared the digestion efficiency by using different buffer conditions, including NaCl/Tris at pH 7.8, NEBuffer^TM^ r1.1 (supplied with RNase 4), and ammonium acetate at pH 5.5 and 7.0 (Figure S4). RNase T1 produced similar cleavage products under all buffer conditions, whereas RNase 4 showed differing fragments, particularly <30 nts, depending on the buffer composition. RNase T1 was active at both pH 5.5 and 7.0, while RNase 4 retained higher activity at pH 7.0. In pure water, RNase 4 showed no activity, and RNase T1 exhibited only minimal activity.

All in all, our investigation indicates that RNase T1 and RNase 4 produce the most variable digestion patterns, while hydrolysates of colicin E5 are homogenous in length and independent of the tested conditions. However, it is important to note that the influence of individual parameters is interdependent; thus, our conclusions are valid only for the tested parameter combinations.

### High resolution MS of hydrolysates by coupling to nanoflow HILIC

Before analysing RNA hydrolysates, we needed an LC-MS/MS method that fulfils our main criteria: (1) utilization of common buffers in liquid chromatography and avoid ion-pairing reagents and (2) ionization which results in good coverage of oligonucleotides and its MS/MS fragments. Commonly, oligonucleotides require the use of ion-pairing reagents to mask their hydrophilicity. As many MS facilities are concerned about ion suppression caused by these reagents, we decided to use an IP-free chromatography and ensure increased accessibility and general applicability for MS core facilities. We decided to use HILIC (hydrophilic liquid interaction chromatography) and exploit the hydrophilicity of oligonucleotides instead. For analytical flow oligonucleotide separation and metabolomics in general, HILIC has been applied by various laboratories (12,22,23). To achieve the highest possible sensitivity, we decided to reduce the flow rate to nanoflow, which has never been achieved for HILIC, yet.

For method development and optimization, we used synthetic RNA and DNA oligonucleotides (sequences in Table S1). Figure 2 shows the mass spectra of a direct injection of a 5-mer, a 20-mer and 30-mer RNA using a HILIC set-up and HILIC-suitable mobile phases and retention times are 38, 55, and 58 minutes, respectively (Figure S5). Oligonucleotides are commonly ionized in negative mode due to the negatively charged phosphodiester backbone (9). Yet, at acidic pH, nucleobases are easily charged and thus analysis in positive mode is also possible (24). We find that both ionization modes are suitable for analysis of oligonucleotides from 5-30 nts (nucleotides), yet the mass spectra differ substantially.

**Figure 2.**
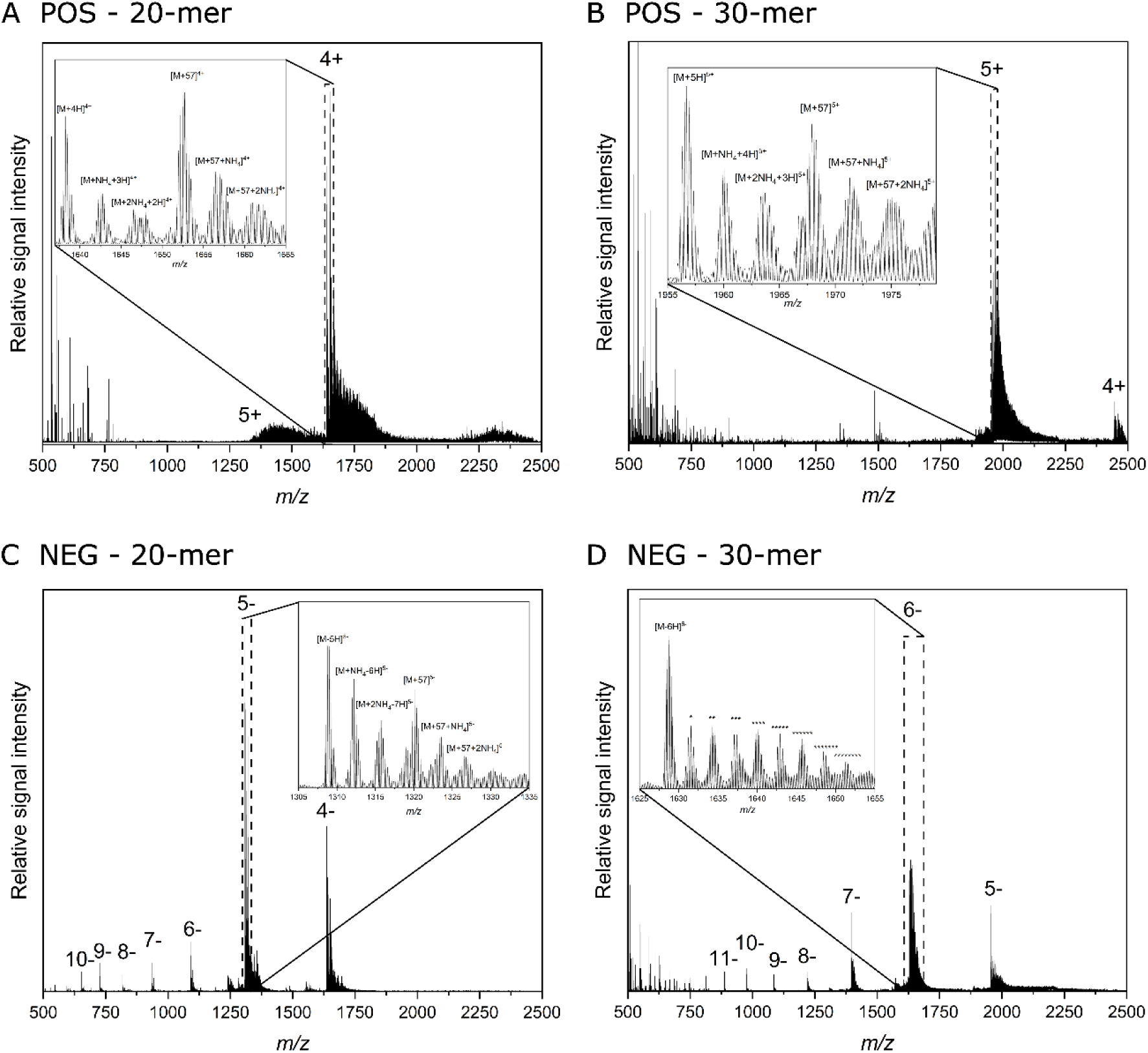
MS^1^ spectra comparison of (**A**) and (**C**) 20-mer (left) and (**B**) and (**D**) 30-mer (right) (synthetic ribonucleotides) between positive ion mode (POS -upper spectra) and negative ion mode (NEG – lower spectra). Charge states of some intense peaks in each spectrum were indicated. A zoom-in spectrum of the most abundant charge states was inserted in each spectrum. The numbers of asterisk (*) indicate the numbers of NH_4_ adducts, e.g. “***” at “6-” indicates [M+3NH_4_-9H]^6-^.

In positive ion mode, we find that all tested oligonucleotides have a dominating charge state in their corresponding mass spectra with signal abundance > 95% (Figure 2 top panel). This contrasts with negative ion mode, where we also find one dominating charge state, but for the longer oligonucleotides additional charge states occur that account for at least 40% of signal abundance (Figure 2 bottom panel). In positive ion mode, the signal of the oligonucleotides is not split into multiple peaks which improves sensitivity compared to negative ionization. Thus, positive ionization is best suited if the goal of analysis is sensitive and targeted analysis of an oligonucleotide. In addition, the spectra are less cluttered in positive ion mode. Negative ionization has the benefit of the higher charge states at MS1 stage, which aids the next step of analysis – MS/MS fragmentation. Oligonucleotides fragment along the phosphodiester and the McCloskey nomenclature defines the possible a,b,c and d-ions (5’-end) and w, x, y and z-ion (3’-end) (Figure S6) (25). The detection of a high number of fragments is needed for correct sequence annotation and supported by high charge states in MS1 and equal distribution of charges at MS/MS. Thus, meaningful MS/MS spectra with a high sequence coverage are ideally received in negative ion mode (Figure S7). Often, MS adducts can disturb the analysis of oligonucleotides by MS. A zoom into the most abundant peak in each spectrum reveals a similar abundance of ammonia adducts in both positive and negative ion mode. This exemplifies the importance of using a clean system with minimal salt load for successful oligonucleotide analysis. Further, we conclude that positive ion mode can be used for targeted analysis of oligonucleotides and negative ion mode for sequencing of oligonucleotides.

### HILIC trap columns are a suitable option for nanoflow HILIC analysis of oligonucleotides

Nanoflow chromatography benefits often from using a trap column as it improves the peak intensity and sensitivity, especially if high loading volumes are needed. To test the applicability of the trap system in the nanoflow HILIC set-up, we installed a HILIC stem trap and loaded synthetic oligonucleotides for 5 minutes at a flowrate of 10 µL/min. The sample was then separated at 250 nL/min using either no analytical column, a reverse phase column or a commercially available HILIC column. We find that the HILIC trap column was able to trap the oligonucleotides, and we received substantial peaks upon elution into the nanoflow path. As expected, the oligonucleotides were not separated when no columns or an RP column were used (Figure S8A/B). In contrast, the oligonucleotides were well separated on the HILIC analytical column (Figure S8C). To further judge the impact of the HILIC trap on the subsequent HILIC separation, the oligonucleotides were directly injected (without a trap column and loading step). In direct injection (Figure S8D), both oligonucleotides elute earlier, but with similar peak shape. Furthermore, we see that the analytical column has a stronger retention than the HILIC trap column, which is important for successful analyte trapping and thus preconcentration. Judging from these observations, we recommend using the trap set-up if volumes exceeding 1 µL need to be injected. For concentrated samples with a 1 µL injection, a direct injection can be used and is in fact recommended as a trap set-up is more prone to leaks and system mal function compared to direct injection.

### High water content disturbs spray and MS detection of oligonucleotides

During analysis of oligonucleotides in both direct injection and trap mode we noticed an MS signal loss once the water content exceeded 50% water. Yet, with the commercial HILIC-amide column, long oligonucleotides were strongly retained, eluted at high water content and their signal was not stable as shown for the 5-, 20- and 30-mer in Figure S9. We found that the spray was more stable at high RF values and for some oligonucleotides the signal could be saved by using an RF of 90 (Figure S9). Yet a loss of spray persisted. If a second nanoflow pump is available, the content of organic solvent can be increased after the chromatographic separation and prior to the ion source. The set-up is shown in Figure S10 and the impact on the MS signal intensity is given in Figure S9C_F. The total ion count (TIC) was stable for all oligonucleotides analysed even at low RF values of 50. In addition, the TIC signal intensity was similar or even better compared to the non-infused samples. A detailed analysis of charge state in MS1 (Figure S11) and fragmentation in MS/MS mode (Figure S12) revealed various positive side effects of the infusion set-up such as the stable spray, high sensitivity and good sequence coverage.

### Lower retention by using the diol column chemistry allows a stable spray

Another option for generation of a continuous and stable spray is the use of another column chemistry, that has a lower retention of oligonucleotides. As nanoflow HILIC columns are not commonly used, we decided to test HILIC materials in self-packed columns. Among the materials tested, the diol chemistry showed the highest usability due to its low interaction with the oligonucleotides while allowing separation. A comparison of the commercial HILIC-amide and the self-packed HILIC-diol column towards the separation of a 20-mer and 30-mer is shown in Figure 3A. As can be seen, the oligonucleotides elute earlier on the diol column and thus at higher organic phase content. Therefore, we did not observe a loss of spray with this column. Regarding the separation ability, we used a mixture of synthetic deoxythymidine oligonucleotides (dT) containing either, 10, 15, 20, 30 or 50 dTs. The separation of these oligonucleotides can be influenced by the steepness of the gradient as seen in Figure 3B. Even at a steep gradient, a resolution above 5 is achieved for all peak pairs (Figure S13), all oligonucleotides elute within 40 minutes and a full MS/MS coverage of the analysed poly-dT oligonucleotides (Figure S14) is obtained.

**Figure 3.**
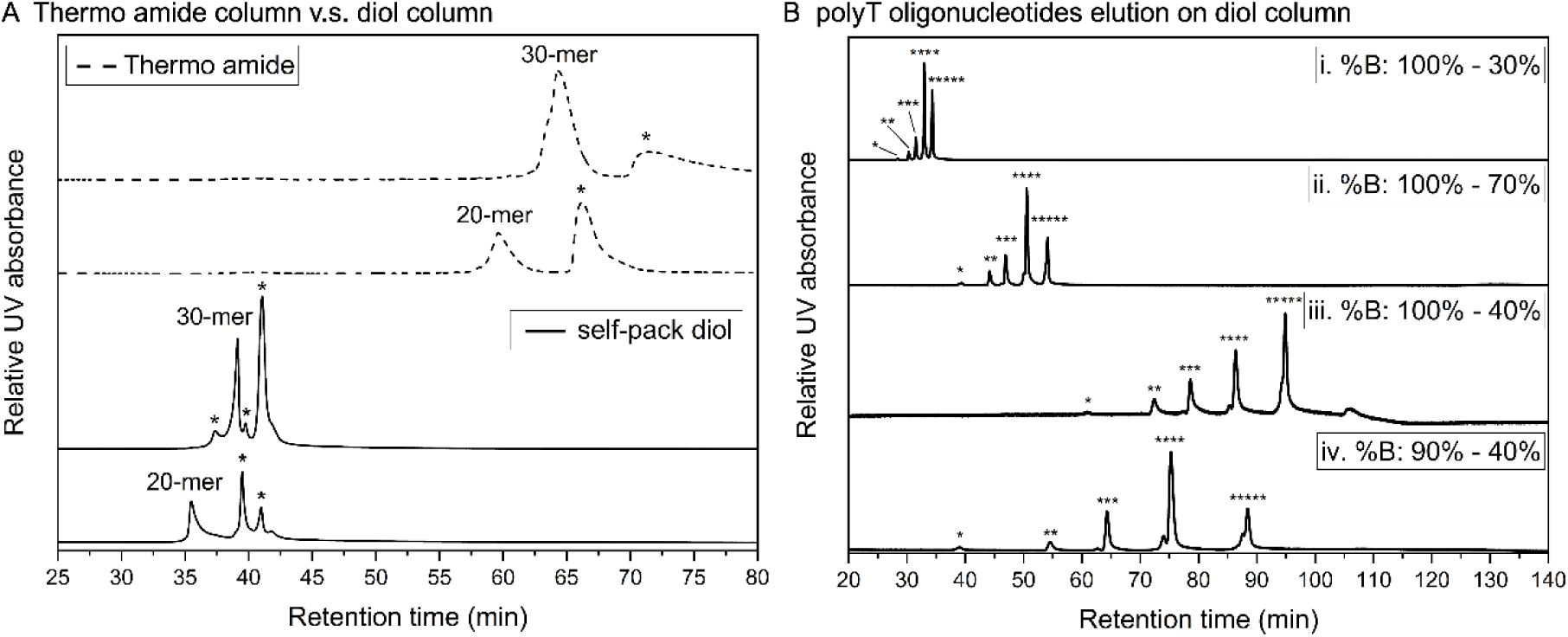
Separation of oligonucleotides on nanoflow HILIC using amide or diol chemistries. (**A**) UV chromatograms of a 20-mer and a 30-mer (synthetic ribonucleotides) eluted on the commercial HILIC-amide column (shown on top in dashed lines) and the self-packed diol column (shown in the bottom in solid lines). Impurities present in these samples were marked by the asterisk (*) signs. Note: The LC gradient used for the self-pack diol column (0.30 μL/min) was slightly (i.e. 9 min) longer than the HILIC-amide column (0.25 μL/min). (**B**) Separation of dT-only containing oligonucleotides with different gradi-ents (I, ii, iii, and vi, which were specified in the supplementary method) on the self-packed diol column. “*”, “**”, “***”, “****”, and “*****” indicated samples “dT10”, “dT15”, “dT20”, “dT30”, and “dT50”, respectively.

Regarding sensitivity, we found that around 0.1 ng/nucleotide, e.g. 2 ng of a 20 nts oligonucleotide, is needed for its detection and successful mapping using the self-packed HILIC-diol nanoflow set-up (Table S3).

### Comparison injection with pure water versus acetonitrile-containing solvent

Since oligonucleotides eluted earlier under the direct injection setup while maintaining acceptable resolution (Figure S8) and because bypassing the trap column simplifies both workflow and troubleshooting, we chose direct injection for the measurement of biological samples. In direct injection, the injection volume (1 µL) exceeds the column dead volume (0.66 µL for a 15 cm × 75 µm column), the solvent composition strongly influences both the consistency of the nano-flow path and the interaction of oligonucleotides with the stationary phase.

Commonly, RNA samples are dissolved in water. To showcase the negative effect of an injection of aqueous RNA into the HILIC system, we first injected a synthetic 8-mer dissolved in 100% water. The oligonucleotide was detected in the breakthrough fraction (Figure S15A), indicating negligible interaction with the stationary phase. In addition, the high-water content disrupted spray stability and caused a pressure increase from ∼60 bar to ∼110 bar (Figure S15A-B). Then the same oligonucleotide was dried and resuspended in 100% mobile phase B, which is the initial condition of our gradient (80% acetonitrile). Under these conditions, the oligonucleotide was retained on the HILIC phase and no pressure spikes or spray instability (Figure S15C-D) were observed. Higher amounts of acetonitrile are not recommended either as RNA precipitates at higher organic content. This phenomenon is constantly used in molecular biology for RNA precipitation using ethanol or acetone and must be considered during RNA sample preparation for HILIC chromatography.

Based on these results, all subsequent injections were performed in 100% mobile phase B to ensure reproducible oligonucleotide separation and stable chromatographic performance.

### Analysis of partial RNA hydrolysates using nano-HILIC-HRMS

After successful detection of synthetic oligonucleotides in our system, we hydrolyzed unmodified tRNA^Ile^ with either RNase T1, RNase 4, or colicin E5. For removal of residual salt and protein, we used the same sample preparation workflow as for our MALDI MS analysis (Zymo spin column) and we found that this method was suitable for good performance of the HILIC nanoflow chromatography. We injected 50 ng of hydrolyzed tRNA onto the column (1µL in direct injection) and the resulting total ion chromatograms are shown in Figure 4. For all three nucleases, cleavage products of reasonable length were found. The oligonucleotides previously detected by MALDI-MS were readily detectable at high resolution and a mass accuracy below 10 ppm. MS/MS spectra of these peaks were also available from these peaks and the observed fragmentation fitted well with the in silico expected fragments (26).

**Figure 4.**
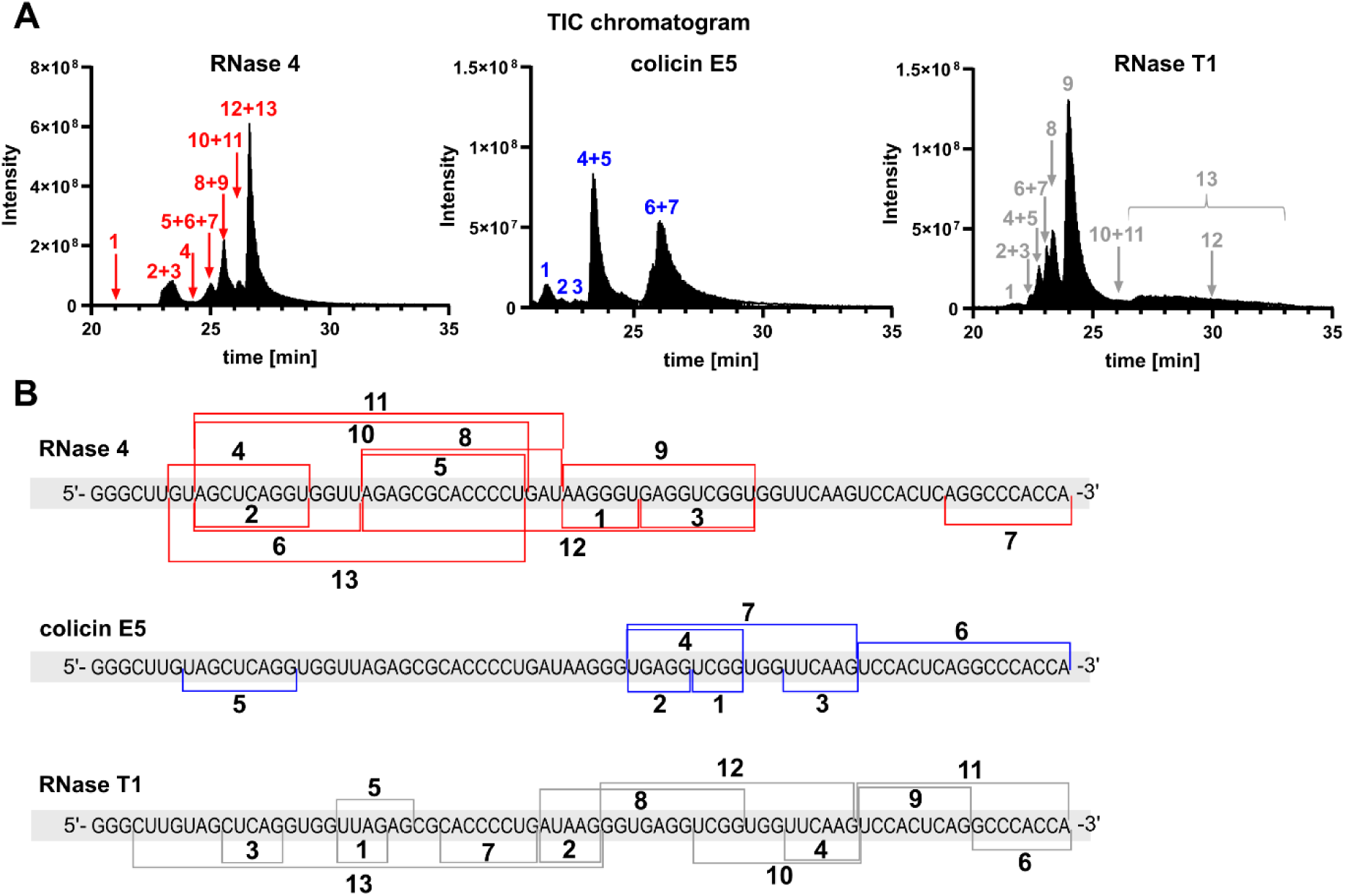
Comparison of RNase T1, RNase 4 and colicin E5 digested unmodified RNA (E. c. tRNA^Ile^_GAU_) in the developed nanoflow-HRMS set-up. (**A**) TIC chromatograms with arrows indicating RT of found fragments. (**B**) Assigned fragments and cleavage sites in the sequence of E. c. tRNA^Ile^_GAU_. Red: RNase 4; blue: colicin E5; grey: RNase T1.

### NASE requires firm knowledge on 3’-end after hydrolysis

With the oligonucleotide-MS method successfully developed, we utilized it to study the substrate specificity of the three RNases in more detail. For product determination and phosphate status determination, we did not utilize a phosphatase at this stage. For this, we wanted to use NASE (NucleicAcidSearchEngine) and its search function for the respective RNase specific cleavage and “unspecific cleavage”. Problematically, when we started analysing our data using the RNase specific cleavage, not all oligonucleotides picked up by MALDI-MS were successfully assigned by NASE. From our previous analysis by MALDI-MS (Figure 1) and manual data analysis (Figure 4) we know that the 3’-end of the oligonucleotides can either carry a 2’,3’-cyclic phosphate, a 3’-phosphate or a 3’-hydroxyl group (Figure 5A-B). We learned that the 3’-end selection is crucial for correct fragment assignment and therefore, NASE was modified to allow selection of different 3’-chemistries.

**Figure 5.**
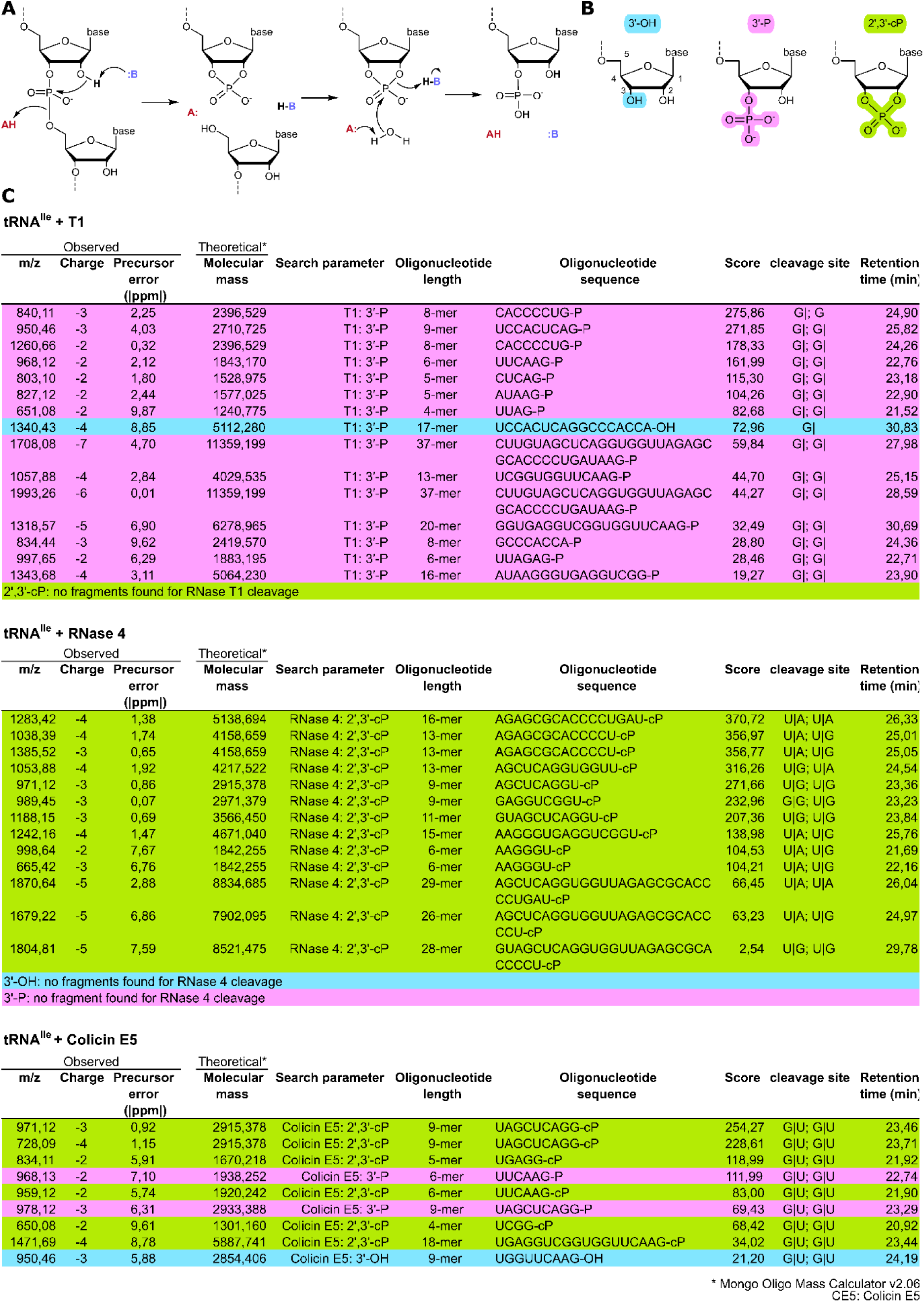
Comparison of RNase 4, colicin E5 and RNase T1 digested unmodified RNA *E. c.* tRNA^Ile^ - GAU) using updated NASE. (**A)** General Ribonuclease-mechanism with AH indicating an acidic amino acid and B indicating a basic amino acid. (**B**) Chemical structures of potential 3’ ends of RNase hydrol-ysis. (**C**) Table of identified fragments of tRNA^Ile^_GAU_ for RNase T1, RNase 4 and colicin E5. cP = cyclic phosphate, P = 3’ phosphate and OH = 3’ hydroxyl. In the column “cleavage site”, the 5’-cleavage site and 3’-cleavage site are separated by semicolon (;).

With the new developments in NASE, we moved forward with analysis of the oligonucleotides produced by the three nucleases. RNase T1 strictly targets G|N sites and consistently produces fragments with 3’-phosphates (Figure 5C). Although full hydrolysis with RNase T1 generates short and less specific products, we achieved longer fragments of up to 39 nt by employing partial digests with reduced enzyme-to-substrate ratios and limited incubation times. This approach provided the highest sequence coverage of all nucleases tested (96.1%), with no fragments detected in the non-specific search mode (Figure S16).

We then moved on to analysis of RNase 4. Using the RNase 4 specific search function in NASE, oligonucleotides emerging from the expected U|A and U|G cleavage were readily detected if combined with 2’,3’-cyclic phosphate search. The fragment lengths ranged from 6 to 29 nucleotides, with sequences such as AGAGCGCACCCCUGAU-cP appearing with high confidence.

In contrast to RNase T1, RNase 4 yielded a narrower fragment distribution, with partial redundancy observed in the middle regions of the tRNA^Ile^ sequence. We searched for 2′,3′-cyclic phosphate, 3′-phosphate and 3′-OH ends in NASE and could only find 2′,3′-cyclic phosphate products, which aligns with the literature of RNase 4 at low concentrations (21). Fragment overlap was most pronounced in uridine-rich loop domains and adjacent stem regions. For example, fragments such as AGAGCGCACCCUGAU-cP (16-mer) and AGAGCGCACCCCU-cP (13-mer) covered common sequence regions but differed in cleavage end points, providing useful redundancy for sequence reconstruction. The unspecific search mode in NASE revealed RNase 4’s capacity to cleave at additional motifs beyond uridine, predominantly only C|A site, as reported (21). For example, detection of AGCUC-cP and AGGU-cP demonstrated cleavage at a single C|A site, either expanding the enzyme’s reported specificity or indicating contamination by an unknown RNase. Other off-target fragments were predominantly 4 to 10 nucleotides in length and consistently showed 2′,3′-cyclic phosphate termini, reinforcing the enzyme’s overall mechanistic consistency. The ability of RNase 4 to cleave at both typical and unexpected sites enabled sequence coverage of 72.7%, of which ∼13% originated from C|A cleavage (Figure S16). Judging from this result, we confirm the suitability of RNase 4 for sequence mapping of highly purified RNAs, where off-target cleavage is a useful feature to gain maximum sequence coverage.

Colicin E5 was also assessed using the G|U-specific cleavage function of NASE. In this targeted search, a high number of cleavage products was detected, with abundant fragments spanning 4 to 18 nucleotides and exhibiting both 2′,3′-cyclic phosphate and 3′-phosphate ends. We found fragments that covered large regions of sequence and often overlapped by one or two nucleotides, a pattern that reflects highly redundant sequence coverage. This overlap was particularly noticeable in U- and G-rich regions, where multiple fragments’ start- and stop-positions created a pattern of quasi-continuous overlap.

Next, we searched for unspecific cleavage sites and found that colicin E5 generated various oligonucleotides outside the expected G|U motif. Detailed review revealed that the non-specificity followed in fact certain rules which indicates an undescribed context-dependent selection of cleavage sites. Cleavage was observed close to GU sites and especially in the context G|G(G)U and GU|C, indicating that colicin E5 slips by 1-2 nucleotides. Furthermore, we observed occurrence of A|A, C|A, C|U, U|C, C|C, G|C, A|C, and U|U which either indicates a broad substrate range or a contamination with an unknown RNase. Those unspecific products have high scores in NASE and make 26% of the total sequence coverage, which was 83.1% (Figure S17).

### RNase 4 and colicin E5 accept major RNA modifications within their substrate sites

Native RNA contains various modified nucleotides. Both RNase 4 and colicin E5 cleave in a uridine-containing dinucleotide context which prompts the question, how the RNases handle the dominant RNA modification pseudouridine (Ψ) within their recognition sequence. To approach this question, tRNA^Ile^ was in vitro transcribed in the presence of ΨTP and no UTP. In the resulting RNA, the HDV ribozyme (Hepatitis-Delta-Virus self-cleaving ribozyme) was unable to release the full lengths tRNA and instead a longer RNA, containing the HDV sequence was observed and purified for further analysis. The fully Ψ-modified RNA and a mixture of regular U-containing tRNA^Ile^ plus cleaved HDV RNA, were incubated with RNase 4 and colicin E5. The resulting fragments were injected into the nanoflow-HILIC-HRMS system, and the total ion count chromatogram is displayed in Figure 6A. Both RNases were active and U-and Ψ-containing fragments were identical in MS1 and MS/MS which clearly proves identical cleavage positions. Ψ-containing fragments eluted slightly later than the corresponding U-containing fragments which is in accordance with expectation due to Ψ’s higher hydrophilicity. From this data we conclude, that (i) RNase 4 and colicin E5 recognize both U and Ψ within their substrate sites and (ii) Ψ-modified oligonucleotides can be distinguished from their unmodified U-containing strands by retention time. Due to Ψ being mass silent, we recommend chemical derivatization for direct detection of Ψ by MS as previously described (27–29).

**Figure 6.**
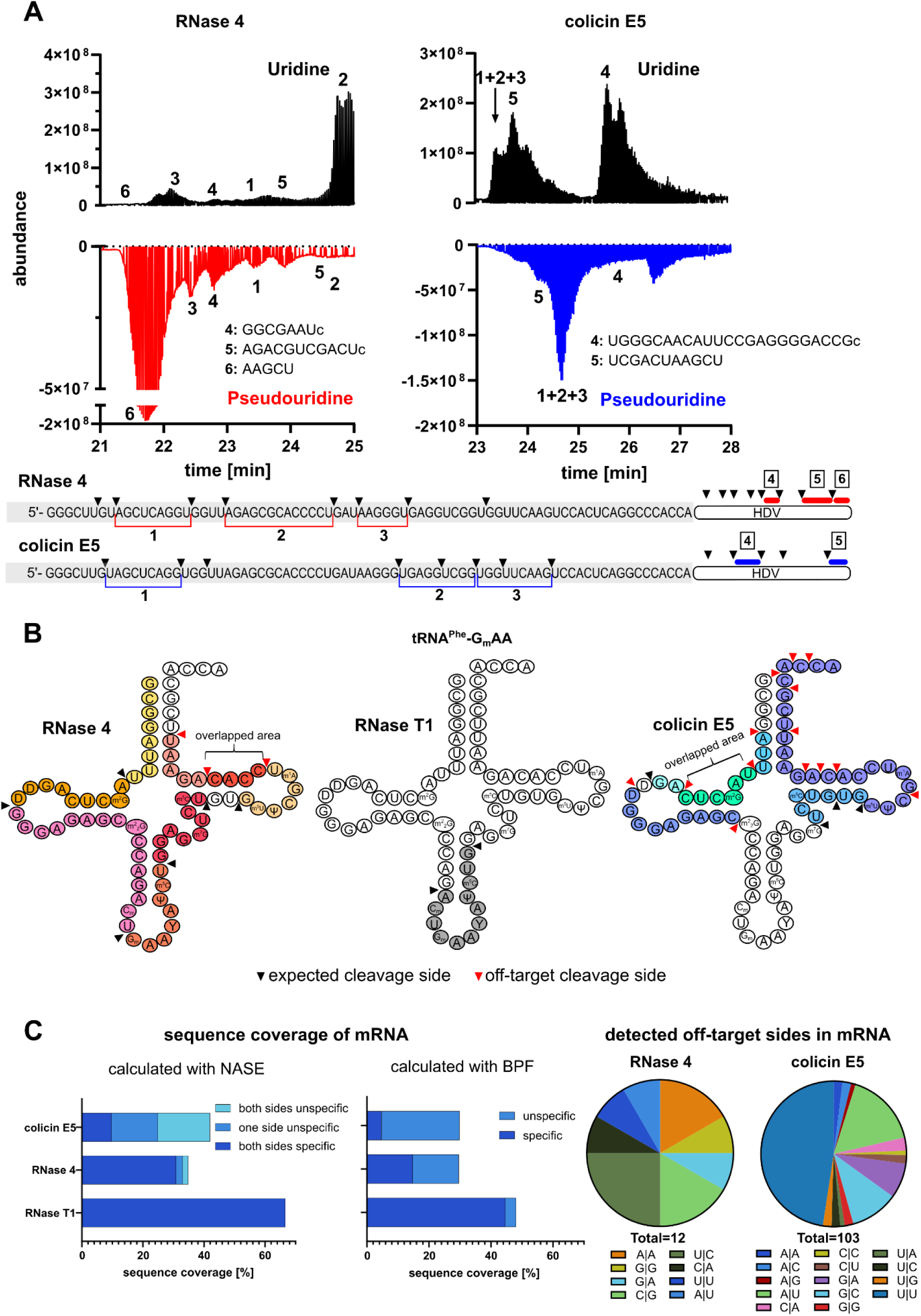
Application of the developed oligonucleotide-MS method with three different RNases to diverse RNA substrates. (**A**) Analysis of a vitro–transcribed (IVT) *E. c.* tRNA^Ile^_GAU_-HDV (Hepatitis-Delta-Virus ribozyme), prepared once with UTP (uridine triphosphate) and once with ΨTP (pseudouridine triphosphate), after digestion with RNase 4 (left) or colicin E5 (right). Top: TIC chromatograms of digested UTP-IVT samples; bottom: TIC chromatograms of ΨTP–IVT samples. Fragments identified by NASE are annotated in the sequence below. The numbered peaks in the chromatograms correspond to the labelled fragments. Red: RNase 4; blue: colicin E5. (**B**) Analysis of native *S. c.* tRNA^Phe^_GmAA_ which is digested with RNase 4, RNase T1 and colicin E5. Detected fragments are highlighted in the cloverleaf secondary structure. Black triangles indicate expected (specific) cleavage sites; red triangles mark off-target cleavage sites. Short sequences detected in two overlapping fragments are labelled as “overlapped area”. (**C**) Analysis of Firefly luciferase (FLuc) mRNA which is digested with RNase 4, RNase T1 and colicin E5. Left: sequence coverage obtained with each RNase, calculated with NASE. Light green: fragments with both termini as expected specific cleavage sites; turquoise: one terminus specific and the other unspecific; dark green: both termini unspecific. Middle: sequence coverage obtained with each RNase, calculated with BPF (ThermoFisher BioPharma Finder). Red: fragments with both termini as expected specific cleavage sites; grey: one or both termini unspecific. Right: distribution of detected off-target cleavage sites in the mRNA for RNase 4 and colicin E5. No off-target sites were detected for RNase T1.

In the next analysis, we challenged our system by using native tRNA^Phe^ from *S. cerevisiae*, that was, until recently, commercially available. For RNase 4, we found successful cleavage for the DΙG and UΙGm, while colicin E5 was active for GΙD, m^7^GΙU, GΙm^5^U and the off-target site CΙm^22^G (Figure 6B). We conclude that both RNases have a low stringency and may accept many U- and G-modifications. Systematic assessment should be the goal of future analysis, albeit the generation of synthetic, modified RNAs remains a major challenge in the field. A more detailed analysis of the tRNA^Phe^ data confirmed our previous observations on site specificity. For RNase 4, 5 out of 7 fragments were from expected target sites, while only 3 out of 16 fragments emerged through the expected substrate cleavage for colicin E5 (Figure S18).

So far, we only studied short tRNAs with a limited availability of the 16 possible dinucleotide contexts. To truly judge the substrate specificity of RNase 4 and colicin 5, we analysed the FLuc-mRNA available from the National Institute of Standardization (NIST, RGTM 10202). For each RNase, 250-400 ng of mRNA hydrolysate were directly injected and separated on the described nanoflow-HILIC-MS set-up. For partial RNase T1 digestion, a 66.6% sequence coverage was determined using NASE and 48.1% using BioPharmaFinder (BPF, proprietary software ThermoFisher). As expected, no off-target cleavage was detected for RNase T1 (Figure 6C). For RNase 4, NASE revealed a 34.8 % and BPF 29.6% sequence coverage and most cleavage sites were the expected UΙG and UΙA sites (Figure 6C) Only 12 detected cleavage sites out of 90 were off-target cleavages with UΙC, CΙG and AΙA dominating.

For colicin E5, NASE found a sequence coverage of 41.9% and BPF of 29.9%. As displayed in Figure 6C, over 100 off-target sites are detectable for colicin E5. Next to GΙU, high cleavage rates are observed for UΙU and AΙU sites. A detailed list of all detected fragments is given in Table S4 and Table S5.

## Discussion

While sequencing-based technologies are ideal to study long RNAs and determine their sequences, they cannot directly identify the chemical nature of modifications. Thus, the accurate analysis of small, highly modified RNAs remains a challenge to the RNA community. LC-MS/MS of RNA by oligonucleotide MS is an orthogonal technology that excels in analysis of small, highly modified RNAs. Currently, oligonucleotide MS is not widely used on native RNAs due to the high amount of RNA needed for analysis. The presented nanoflow-HILIC chromatography enables a highly sensitive analysis of sub-microgram amounts of RNA, which in turn enables the modification-aware analysis of small, densely modified RNAs but also long, enriched RNAs. For native tRNA, an injection of 25-50 ng is sufficient for complete sequence and modification coverage. For mRNA, 250-400 ng injections are sufficient for high sequence coverage of an 1850 nts long mRNA. On average, 20-100-fold less RNA is needed for oligonucleotide MS using nanoflow HILIC compared to analytical flow methods (30). This will be especially interesting for the analysis of native small RNAs, such as tRNA, tiRNA, and miRNAs that are highly modified.

Yet, the sensitivity of nanoflow chromatography has a high price, namely low system robustness and unexpected system failures due to leaks and spray instabilities. Thus, we recommend the system mainly for analyses where extremely high sensitivity is needed.

Regarding the substrate specificity, we find that RNase T1 remains to be the most reliable RNase in terms of substrate specificity. If the ratio of RNA and RNase is well dosed, partial cleavage occurs and oligonucleotides with unique identity emerge. This is particularly useful for approaches with heterogenous RNA mixtures. Here, in silico digestion is simplified by the combination of G-specificity and missed-cleavage calculation. For single RNA analysis, both RNase 4 and colicin E5 are highly useful (Figure S19). Regarding the analysis of RNA mixtures, we assume that the off-target cleavage poses a major challenge, especially with colicin E5. Off-target cleavage increases the number of emerging oligonucleotides from each RNA within the mixture; cluttered chromatograms and MS spectra may be a consequence. While “decluttering” is possible by enhancing chromatographic separation even further and making MS analysis faster (or using data independent MS/MS analysis), data analysis of the multiple fragments remains a key bottleneck. RNase 4 showed some off-target cleavage as well, but substantially less than colicin E5. Thus, both RNases need to be further tested, and cleavage product formation must be systematically assessed using modified RNAs. In this work, we deliberately avoided the use of phosphatase with the goal to determine the 3’-chemistry of the hydrolysis product. From our analyses (Figure 5), we now know that both RNase T1 and RNase 4 produce uniform 3’-ends (3’-phosphate and cyclic phosphate, respectively) which results in the formation of defined oligonucleotides per cleavage. For colicin E5, we see more variety in the 3’-products and thus the use of alkaline phosphatase (24) for generation of uniform 3’-hydroxylated oligonucleotides may be useful.

With a highly sensitive method and well-defined RNases now in hand, many hurdles towards the use of oligonucleotide MS in routine analysis of native RNAs and cellular lysates have been overcome. A remaining hurdle is to make the technology quantitative. For this, we can borrow concepts already established for protein analysis. One exploits the chemical reactivity of amino acids by adding an isobaric tag to each peptide and the second utilizes metabolic labelling of proteins using stable isotopes (SILAC). An isobaric tag is not yet described for oligonucleotides but would be the ideal solution. A SILAC-comparable technology for RNA exists in human cell culture, yeast and bacteria and is called NAIL (nucleic acid isotope labelling) (31). With NAIL, Carbon-13 and Nitrogen-15 isotopologues of oligonucleotides emerge and can be used as spike-in reference standards. Although feasible in the wet lab, data analysis of NAIL-derived RNA is not yet possible. Here, further developments are needed in both NASE and BPF. With our developments and those from colleagues in the field, we foresee that the next 3-5 years will give rise to an oligonucleotide MS method of sufficient sensitivity which is widely accepted in MS core facilities to allow quantitative and sequence-aware analysis of epitranscriptomic changes in human cells.

## DATA AVAILABILITY

*The oligonucleotide MS data underlying this article have been deposited to the Proteo-meXchange Consortium via the PRIDE* (*32*) *partner repository with the dataset identifier PXD070755.* [**Project accession:** PXD070755 and PXD070794 **Project DOI:** N/A; Reviewer account details: **Username:****Password:** Sz0UmMmFaZgB].

## SUPPLEMENTARY DATA

Supplementary Data are available at NAR online.

## AUTHOR CONTRIBUTIONS

Yuyang Qi: Conceptualization, Formal analysis, Data curation, Methodology, Validation, Writing—original draft. Chengkang Li: Conceptualization, Formal analysis, Data curation, Investigation, Methodology, Writing—original draft. Nur Yesiltac-Tosun: Conceptualization, Formal analysis, Data curation, Methodology, Writing—original draft. Jannick Schicktanz: Formal analysis, Data curation. Leona Rusling: Formal analysis, Data curation, Writing—original draft. Steffen Kaiser: Formal analysis, Data curation. Sam Wein: Methodology (NASE). Stefanie Kaiser: Conceptualization, Formal analysis, Visualization, Writing—original draft, review & editing, Acquisition of funding.

## ACKNOWLEDGEMENTS

Stefanie Kaiser and Sam Wein are members of the Human RNome Project Consortium.

## LITERATURE

1. Cappannini, A., Ray, A., Purta, E., Mukherjee, S., Boccaletto, P., Moafinejad, S.N., Lechner, A., Barchet, C., Klaholz, B.P., Stefaniak, F., et al. (2024) MODOMICS: a database of RNA modifications and related information. 2023 update. Nucleic Acids Res, 52, D239–D244.

2. Suzuki, T. (2021) The expanding world of tRNA modifications and their disease relevance. Nature Reviews Molecular Cell Biology, 22, 375–392.

3. Kariko, K., Muramatsu, H., Welsh, F.A., Ludwig, J., Kato, H., Akira, S. and Weissman, D. (2008) Incorporation of pseudouridine into mRNA yields superior nonimmunogenic vector with increased translational capacity and biological stability. Mol Ther, 16, 1833–1840.

4. Yu, Q., Liu, S., Yu, L., Xiao, Y., Zhang, S., Wang, X., Xu, Y., Yu, H., Li, Y., Yang, J., et al. (2021) RNA demethylation increases the yield and biomass of rice and potato plants in field trials. Nature Biotechnology, 39, 1581–1588.

5. Adams, M.D., Boileau, E., Bujnicki, J.M., Cheung, V.G., Conticello, S.G., Dedon, P., Dieterich, C., Gallo, A., Göke, J., Helm, M., et al. (2025) Unlocking the regulatory code of RNA: launching the Human RNome Project. Genome Biology, 26, 367.

6. Katopodi, X.-L., Begik, O. and Novoa, Eva M. (2025) Toward the use of nanopore RNA sequencing technologies in the clinic: challenges and opportunities. Nucleic Acids Research, 53.

7. Jenjaroenpun, P., Wongsurawat, T., Wadley, T.D., Wassenaar, T.M., Liu, J., Dai, Q., Wan-chai, V., Akel, N.S., Jamshidi-Parsian, A., Franco, A.T., et al. (2021) Decoding the epitranscriptional landscape from native RNA sequences. Nucleic Acids Res, 49, e7.

8. Yoluc, Y., Ammann, G., Barraud, P., Jora, M., Limbach, P.A., Motorin, Y., Marchand, V., Tisne, C., Borland, K. and Kellner, S. (2021) Instrumental analysis of RNA modifications. Crit Rev Biochem Mol Biol, 56, 178–204.

9. Limbach, P.A., Crain, P.F. and McCloskey, J.A. (1995) Characterization of oligonucleotides and nucleic acids by mass spectrometry. Curr Opin Biotechnol, 6, 96–102.

10. Jiang, T., Yu, N., Kim, J., Murgo, J.R., Kissai, M., Ravichandran, K., Miracco, E.J., Presnyak, V. and Hua, S. (2019) Oligonucleotide Sequence Mapping of Large Therapeutic mRNAs via Parallel Ribonuclease Digestions and LC-MS/MS. Anal Chem, 91, 8500–8506.

11. Jora, M., Lobue, P.A., Ross, R.L., Williams, B. and Addepalli, B. (2019) Detection of ribonucleoside modifications by liquid chromatography coupled with mass spectrometry. Biochim Biophys Acta Gene Regul Mech, 1862, 280–290.

12. Lobue, P.A., Yu, N., Jora, M., Abernathy, S. and Limbach, P.A. (2019) Improved application of RNAModMapper - An RNA modification mapping software tool - For analysis of liquid chromatography tandem mass spectrometry (LC-MS/MS) data. Methods, 156, 128–138.

13. Thakur, P., Estevez, M., Lobue, P.A., Limbach, P.A. and Addepalli, B. (2020) Improved RNA modification mapping of cellular non-coding RNAs using C- and U-specific RNases. Analyst, 145, 816–827.

14. Holvec, S., Barchet, C., Lechner, A., Fréchin, L., De Silva, S.N.T., Hazemann, I., Wolff, P., von Loeffelholz, O. and Klaholz, B.P. (2024) The structure of the human 80S ribosome at 1.9 Å resolution reveals the molecular role of chemical modifications and ions in RNA. Nature Structural & Molecular Biology, 31, 1251–1264.

15. Wein, S., Andrews, B., Sachsenberg, T., Santos-Rosa, H., Kohlbacher, O., Kouzarides, T., Garcia, B.A. and Weisser, H. (2020) A computational platform for high-throughput analysis of RNA sequences and modifications by mass spectrometry. Nat Commun, 11, 926.

16. Wolf, E.J., Grunberg, S., Dai, N., Chen, T.H., Roy, B., Yigit, E. and Correa, I.R. (2022) Human RNase 4 improves mRNA sequence characterization by LC-MS/MS. Nucleic Acids Res, 50, e106.

17. Pfeuffer, J., Sachsenberg, T., Alka, O., Walzer, M., Fillbrunn, A., Nilse, L., Schilling, O., Reinert, K. and Kohlbacher, O. (2017) OpenMS - A platform for reproducible analysis of mass spectrometry data. J Biotechnol, 261, 142–148.

18. Rost, H.L., Sachsenberg, T., Aiche, S., Bielow, C., Weisser, H., Aicheler, F., Andreotti, S., Ehrlich, H.C., Gutenbrunner, P., Kenar, E., et al. (2016) OpenMS: a flexible open-source software platform for mass spectrometry data analysis. Nat Methods, 13, 741–748.

19. Schilling, B., MacLean, B.X., D’Souza, A., Rardin, M.J., Shulman, N.J., MacCoss, M.J. and Gibson, B.W. (2014) In Eyers, C. E. and Gaskell, S. (eds.), Quantitative Proteomics. The Royal Society of Chemistry, pp. 0.

20. Thompson, J.E., Venegas, F.D. and Raines, R.T. (1994) Energetics of catalysis by ribonucleases: fate of the 2’,3’-cyclic phosphodiester intermediate. Biochemistry, 33, 7408–7414.

21. Wolf, E.J., Grünberg, S., Dai, N., Chen, T.H., Roy, B., Yigit, E. and Corrêa, I.R. (2022) Human RNase 4 improves mRNA sequence characterization by LC-MS/MS. Nucleic Acids Res, 50, e106.

22. Huang, M., Xu, X., Qiu, H. and Li, N. (2021) Analytical characterization of DNA and RNA oligonucleotides by hydrophilic interaction liquid chromatography-tandem mass spectrometry. J Chromatogr A, 1648, 462184.

23. Tang, D.Q., Zou, L., Yin, X.X. and Ong, C.N. (2016) HILIC-MS for metabolomics: An attractive and complementary approach to RPLC-MS. Mass Spectrom Rev, 35, 574–600.

24. Hagelskamp, F., Borland, K., Ramos, J., Hendrick, A.G., Fu, D. and Kellner, S. (2020) Broadly applicable oligonucleotide mass spectrometry for the analysis of RNA writers and erasers in vitro. Nucleic Acids Res, 48, e41.

25. McLuckey, S.A., Van Berkel, G.J. and Glish, G.L. (1992) Tandem mass spectrometry of small, multiply charged oligonucleotides. J Am Soc Mass Spectrom, 3, 60–70.

26. Rozenski, J. (1999).

27. Carlile, T.M., Rojas-Duran, M.F., Zinshteyn, B., Shin, H., Bartoli, K.M. and Gilbert, W.V. (2014) Pseudouridine profiling reveals regulated mRNA pseudouridylation in yeast and human cells. Nature, 515, 143–146.

28. Schwartz, S., Bernstein, D.A., Mumbach, M.R., Jovanovic, M., Herbst, R.H., Leon-Ricardo, B.X., Engreitz, J.M., Guttman, M., Satija, R., Lander, E.S., et al. (2014) Transcriptome-wide mapping reveals widespread dynamic-regulated pseudouridylation of ncRNA and mRNA. Cell, 159, 148–162.

29. Hermon, S.J., Sennikova, A. and Becker, S. (2024) Quantitative detection of pseudouridine in RNA by mass spectrometry. Sci Rep, 14, 27564.

30. Vanhinsbergh, C.J., Criscuolo, A., Sutton, J.N., Murphy, K., Williamson, A.J.K., Cook, K. and Dickman, M.J. (2022) Characterization and Sequence Mapping of Large RNA and mRNA Therapeutics Using Mass Spectrometry. Anal Chem, 94, 7339–7349.

31. Heiss, M., Hagelskamp, F., Marchand, V., Motorin, Y. and Kellner, S. (2021) Cell culture NAIL-MS allows insight into human tRNA and rRNA modification dynamics in vivo. Nat Commun, 12, 389.

32. Perez-Riverol, Y., Bandla, C., Kundu, D.J., Kamatchinathan, S., Bai, J., Hewapathirana, S., John, N.S., Prakash, A., Walzer, M., Wang, S., et al. (2025) The PRIDE database at 20 years: 2025 update. Nucleic Acids Res, 53, D543–D553.

