## Supplement Information for "Ion-Pair–Free Nanoflow HILIC–MS With RNase Benchmarking for Native RNA"

§ contributed equally to the manuscript

**Table S1** RNA and DNA used in this manuscript

| name | sequence | supplier |
|---|---|---|
| 5 mer | AUUUG | IBA |
| 8 mer | UUUCCCCG | IBA |
| 9 mer | GUUUCCGUA | Sigma |
| 10 mer | AAAUCCAUUG | IBA |
| 13 mer | GGCGGAAACACCA | Sigma |
| 20 mer | UGAGGCAGGAGGUUGAAUAG | Dharmacon |
| 30 mer | AAGCUGAGGCAGGAGGUUGAAUAGCAUGCA | Dharmacon |
| 40 mer | GUA GUC GUG GCC GAG UGG UUA AGG CGA UGG ACU UGA AAU C | IBA |
| DNA T10 | TTTTTTTTTT | Sigma |
| DNA T15 | TTTTTTTTTTTTTTT | Sigma |
| DNA T20 | TTTTTTTTTTTTTTTTTTTT | Sigma |
| DNA T30 | TTTTTTTTTTTTTTTTTTTTTTTTTTTTTT | Sigma |
| DNA T50 | TTTTTTTTTTTTTTTTTTTTTTTTTTTTTTTTTTTTTTTTTTTTTTTTTT | Sigma |
| 77 mer tRNA$^{Ile}$ | GGGCUUGUAGCUCAGGUGGUUAGAGCGCACCCCUGAUAAGGGUGA GGUCGGUGGUUCAAGUCCACUCAGGCCCACCA | IVT (in house) |

**Table S2** defined parameter for RNase T1, RNase 4 and colicin E5 for the test of different reaction conditions

| Nuclease | Enzyme to substrate ratio | Temperature [°C] | Incubation time [min] | buffer |
|---|---|---|---|---|
| RNase T1 | 10 U/µg tRNA | 37 | 30 | 90 mM NaCl + 25 mM Tris HCl pH 7.5 |
| RNase 4 | 15 U/µg tRNA | 37 | 60 | NEBuffer™ r1.1* |
| Colicin E5 | 4 mol Colicin E5 to 1 mol substrate | 37 | 30 | 90 mM NaCl + 25 mM Tris HCl pH 7.5 |

*contains 10 mM Bis-Tris-Propane-HCl, 10 mM $MgCl_2$, 100 µg/ml Recombinant Albumin, pH 7.0@25°C

**Table S3.** Calculated limit of detection (LoD) and limited of quantification (LoD).*

| | dT10 | dT15 | dT20 | dT30 | dT50 |
|---|---|---|---|---|---|
| LoD (pmol) | 0.21 | 0.39 | 0.23 | 0.40 | 0.27 |
| LoQ (pmol) | 0.63 | 1.19 | 0.70 | 1.22 | 0.81 |
| LoD (fmol/nt) | 20.78 | 26.21 | 11.54 | 13.46 | 5.36 |
| LoQ (fmol/nt) | 62.98 | 79.43 | 34.98 | 40.80 | 16.24 |
| LoD (ng/nt) | 0.06 | 0.12 | 0.07 | 0.12 | 0.08 |
| LoQ (ng/nt) | 0.19 | 0.36 | 0.21 | 0.37 | 0.25 |

*Calibration curves were prepared using mixtures of DNA oligonucleotides, including dT10, dT15, dT20, dT30, and dT50, in different injection amounts, i.e. 0.2, 0.4, 1, 2, and 4 pmol.

**Table S4.** Detected cleavage fragments from FLuc mRNA, calculated with NASE. Fragments with * are unspecific uncleavage products.

| RNase T1 | colicin E5 | RNase 4 |
|---|---|---|
| AAAAAGp | UAACAACCGCGAAAAAGc | AAAACGGAUUc |
| AAAAAGUUGCGCGGAGp | UACCAGAGc | AAAGAAAGGCCCGGCGCCAUUCUc |
| AAAAAUCAGp | UACCCCCGc | AACAACCGCGAAAAAGUUc |
| AAAACGp | UACCGAAAGGc | AAGAAGAAAUc |
| AAAACUCGp | UAUGCAGc | AAUACGACUCACUAUc |
| AAAACUCUCUUCAAUUCUUUAUGp | UCACAUCUCAUCUACCUCCCGGc | AAUCCAGAAAAUUc |
| AAAAGp | UCGCCAGc | ACAAAAUUCAAAGUc |
| AAAAUUAUUAUCAUGp | UCGGGGAAGCGGc | ACAAGCGAUc |
| AAAGp | UCUUCCCGACGAUGACGCCGGc | ACACCCGAGGGGGAUc |
| AAAUAUAAGp | UGAUAAUAGGCUGGAGCCUCGGc | ACACGAAAUUc |
| AACAAUUGp | UGGACAUCACGc | ACACUCGGAUc |
| AACACUUCUUCAUAGp | UGGACGAAGc | ACCAACCCUc |
| AACGp | UGGAUUACGc | ACCCCCGUc |
| AACUCCUCUGp | UGGCAGGc | ACCGAAAGGUCUUc |
| AACUUCCCGp | *AAAAAAAAUc | ACCGGAUACCUc |
| AAGGGCGGAAAGAUCGp | *AAAACGCUc | ACCGGGAAAACGCUc |
| AAUAAAGp | *AAAUACGAc | ACCUCCCGGUUUUc |
| AAUACAAAUCACAGp | *AAAUACGAUc | ACGACAAGGAUc |
| AAUACUUCGp | *AACCCUAUc | ACGAUCCCUUCAGGAUUc |
| AAUAUGp | *AAGAAAAAUCc | ACGCGGAAUc |
| AAUUAUGp | *ACAAAAAUc | AGGCUGGAGCCUCGGUc |
| ACAAAACAAUUGp | *ACGAAAUc | GAACUCCUCUc |
| ACAAAUACGp | *ACUCUGAUc | GAAGAGGAGCUc |
| ACAAGp | *AGAAUCGc | GAAGCGAAGGUUc |
| ACAUCACGp | *AUCAAAAUc | GACAAAACAAUUc |
| ACAUUUAUAAUGp | *CAAAAAAAAc | GACGCCGGUc |
| ACCAACGp | *CAAAAAAc | GACGGAAAAAGAGAUCGUc |
| ACCUAUGp | *CAAAAAAUc | GAGACUACAUCAGCUc |
| ACUACAUCAGp | *CAAAAAAUUc | GCAAAAAAAAUUc |
| AUAAACCGp | *CAAAACGCUc | GCAAAAAAUUUUc |
| AUAACGp | *CACCCGc | GCACCCGUc |
| AUAAUAGp | *CACCUCUc | GCCAGAGAUCCUc |
| AUAAUGp | *CACUAUAGGGc | GCGUCAGAUUCUCGCAUc |
| AUACCAGp | *CACUGAUAAc | GGAACCGCUc |
| AUACGp | *CAGCCUACCGc | GGAAGACGCCAAAAACAUc |
| AUACUGp | *CAUAGAACUGc | GGACAUCACGUc |
| AUAGp | *CAUCAGGAGc | GGAGAGCAACUc |
| AUAUGp | *CAUCAGGAGUc | GGAGCCUCGGUGGCCUc |
| AUAUUGp | *CAUCGAGc | GGCAGAAGCUc |
| AUAUUUGp | *CGAAAAAGc | GGCCCCCGCUc |
| AUCCCUUCAGp | *CGGAAUACUc | GGCCCUUCCGCAUc |
| AUCCUAUUUUUGp | *CUCAACAGc | GGGCAUUUCGCAGCCUc |

AUCCUCAUAAAGp
AUCUACUGp
AUCUUCACCUAGp
AUUACAAAAUUCAAAGp
AUUACACCCGp
AUUACCAGp
AUUAUGp
AUUCUAAAACGp
AUUCUCGp
AUUUAUCUAAUUUACACGp
AUUUCAGp
AUUUCGp
AUUUUAAGp
AUUUUGp
CAAAAAAAAUUACCAAUAAUCCAGp
CAAAAAAUUUUGp
CAAAACGp
CAACGp
CAACUGp
CAAUCAAAUCAUUCCGp
CACAUAUCGp
CACCCGp
CACCCGUACCCCCGp
CACCUCUUUCGp
CACUCUGp
CACUGp
CAGCAGp
CAGUGGp
CAUAAGp
CAUAGp
CAUCGp
CAUGp
CCAAAAACAUAAAGp
CCAAAAGp
CCAAGp
CCACCAUGp
CCAUUCUAUCCGp
CCCCCCUGp
CCCCCGp
CCCCUUGp
CCCUUCCGCAUAGp
CCCUUCCGp
CCUACCGp
CCUCGp
CCUUGp
CGCCAUUCUAUCCGp
CUACAGp
CUACAUUCUGp
CUAUGp
CUAUUCUGp
CUCAACAGp
CUCACUGp
CUUACUGp
CUUCCAUCUUCCAGp
CUUCUGp
CUUCUUGp
CUUUUACAGp
GAUUAGCAGAGCGAGGp
GAUUAGCAGAGCGAGGUAUGp
UAAAAAGp
UAAACAAUCCGp
UAAAGp
*CUCAUCUc
*GAAAGAAGc
*GUCAAAAAUAc
*UAAAGc
*UAAGAAGAAAc
*UAAUGAACGc
*UACAAAAc
*UACAAAAUc
*UACACGAAAc
*UACACGAAAUc
*UACACGc
*UACAGAUGc
*UACCAACCCUAc
*UACCAAUAAc
*UACCAGGGAc
*UACCAGGGAUc
*UACCUAAGGGc
*UACGAUCCCUc
*UACGCCCUGGc
*UACUGCGAUc
*UACUUCGAAAc
*UAUAAUGAACGc
*UAUAGAUc
*UAUCAUGGAc
*UAUCAUGGAUc
*UAUCGGAGc
*UAUCUAAUc
*UAUGCCGGc
*UAUGGGCAUc
*UCAAAGc
*UCAAAUCAc
*UCAAAUCAUc
*UCAAUUCUc
*UCACAGAAUCGc
*UCACAUCUCAc
*UCACAUCUCc
*UCACGUUAc
*UCAUGGAUc
*UCCAUCACGc
*UCCAUCACGGc
*UCCGGc
*UCCUGCACCCGc
*UCCUGGAACAAc
*UCGAAAGAAGc
*UCGAGc
*UCGAUAUc
*UCGGAGc
*UCUAAAACGGAc
*UCUCGc
*UCUCGCAc
*UCUCGCAUGc
*UCUGAUc
*UCUUCAUAGc
*UGAACGc
*UGAACUc
*UGAACUUCCc
*UGAAGCGAAGc
*UGAAGCGAAGGc
*UGAAUAAAGc
*UGAAUACGAc
*UGAAUACGAUc
*UGACAAAACAAc
GGGCGCGUUc
GGGCUCACUc
GGGCUCACUGAGACUc
GGUUCCUGGAACAAUUc
GUCAGAGGACCUc
GUUCCAUUCCAUCACGGUUUUGGAAUc
*CAUUCCGGAUc
*CUUCCAGGGAUc
*GGGCGGCc
*ACCGGAAAACUc
*CGACGCAAGc
*GGGCGCGc
*UUUUGGCAAc

UAACAACCGp
UAACUAUCGp
UACACGp
UACCAACCCUAUUUUCAUUCUUCGp
UACCCCCGp
UACGp
UAUGp
UCACAUCUCAUCUACCUCCCGp
UCCUUUGp
UCUUAAUGp
UCUUACCGp
UCUUCCCGp
UCUUUAAUUAAAUACAAAGp
UCUUUGp
UUAAUCAGp
UUACAACACCCCAACAUCUUCGp
UUACCUAAGp
UUAUGp
UUAUUGp
UUAUUUAUCGp
UUCCAUAGp
UUCCAUUCCAUCACGp
UUCCAUUUUUUGp
UUCCGACCCUGCCGp
UUCCUGp
UUUACUACACUCGp
UUUCCAAAAAGp
UUUCCCCCUGGAAGCUCCCUCGp
UUUCCCGp
UUUUAAUGp
UUUUGp
UUUUUACGp

*UGACAAAUc
*UGACAAGGAc
*UGACCGCUc
*UGAUAc
*UGAUCGc
*UGCAAAAAAAAc
*UGCAAAACGCUc
*UGCAGc
*UGCCCCUc
*UGCUAGc
*UGCUCAACAGc
*UGGAAc
*UGGAAUGc
*UGGAGc
*UGGAUUCUAAAc
*UGGCAAUCc
*UGGCCCUc
*UGGCCUAGCUc
*UGGGCAUc
*UGGGCGc
*UGGGCGCGc
*UGGGCGGCc

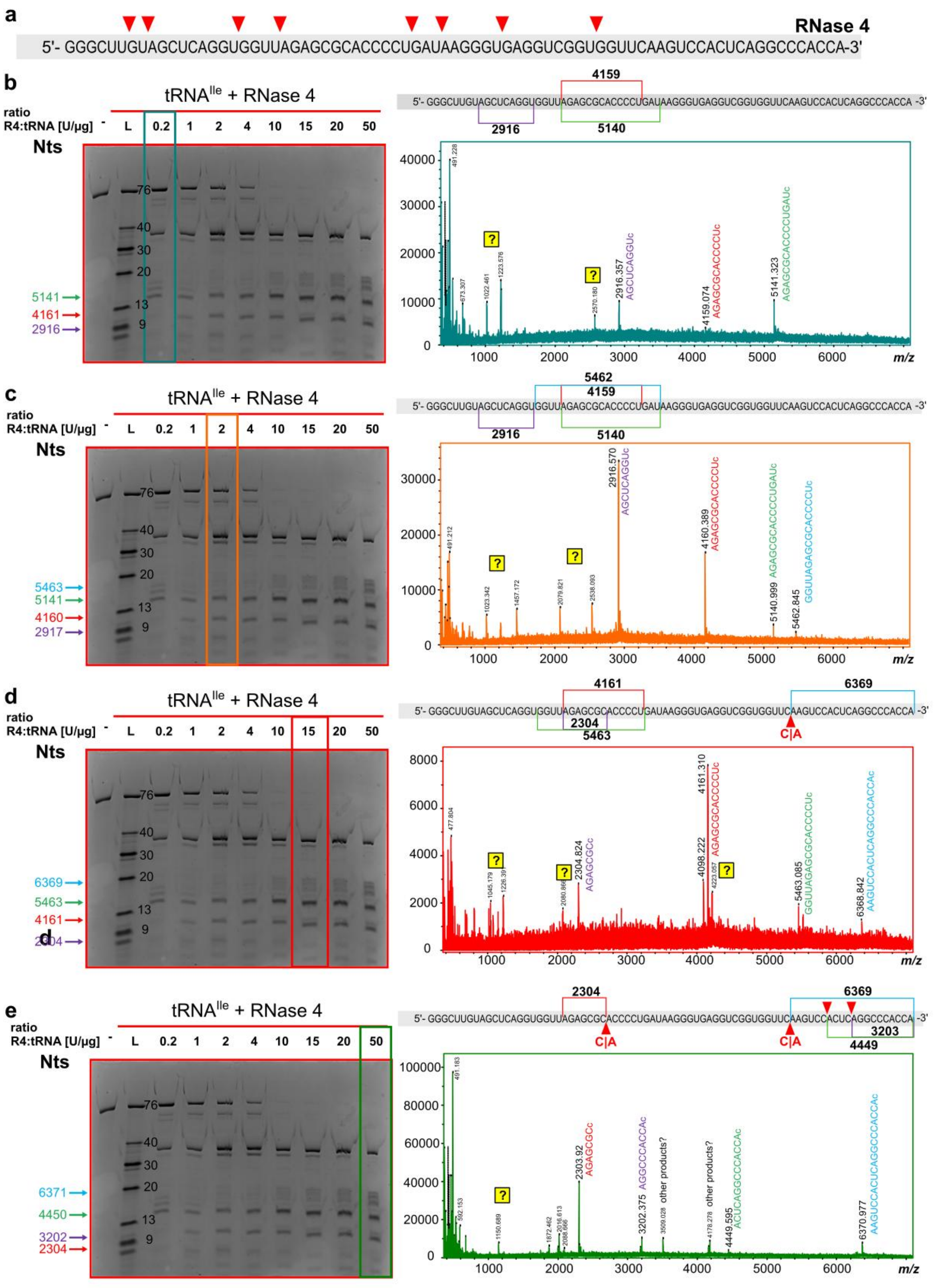

**Figure S1**: Substrate specificity of RNase 4 at different enzyme to substrate ratios. a) sequence of an unmodified tRNA (E. c. tRNA-Ile-GAU) and the expected cleavage sites of RNase 4 hydrolysis. b-e) polyacrylamide gel of RNase 4 digested E. c. tRNA-Ile-GAU with different enzyme to substrate ratios and the corresponding MALDI-MS spectra. Assigned fragments and cleavage sites are added to the MS spectrum and in the sequence above.

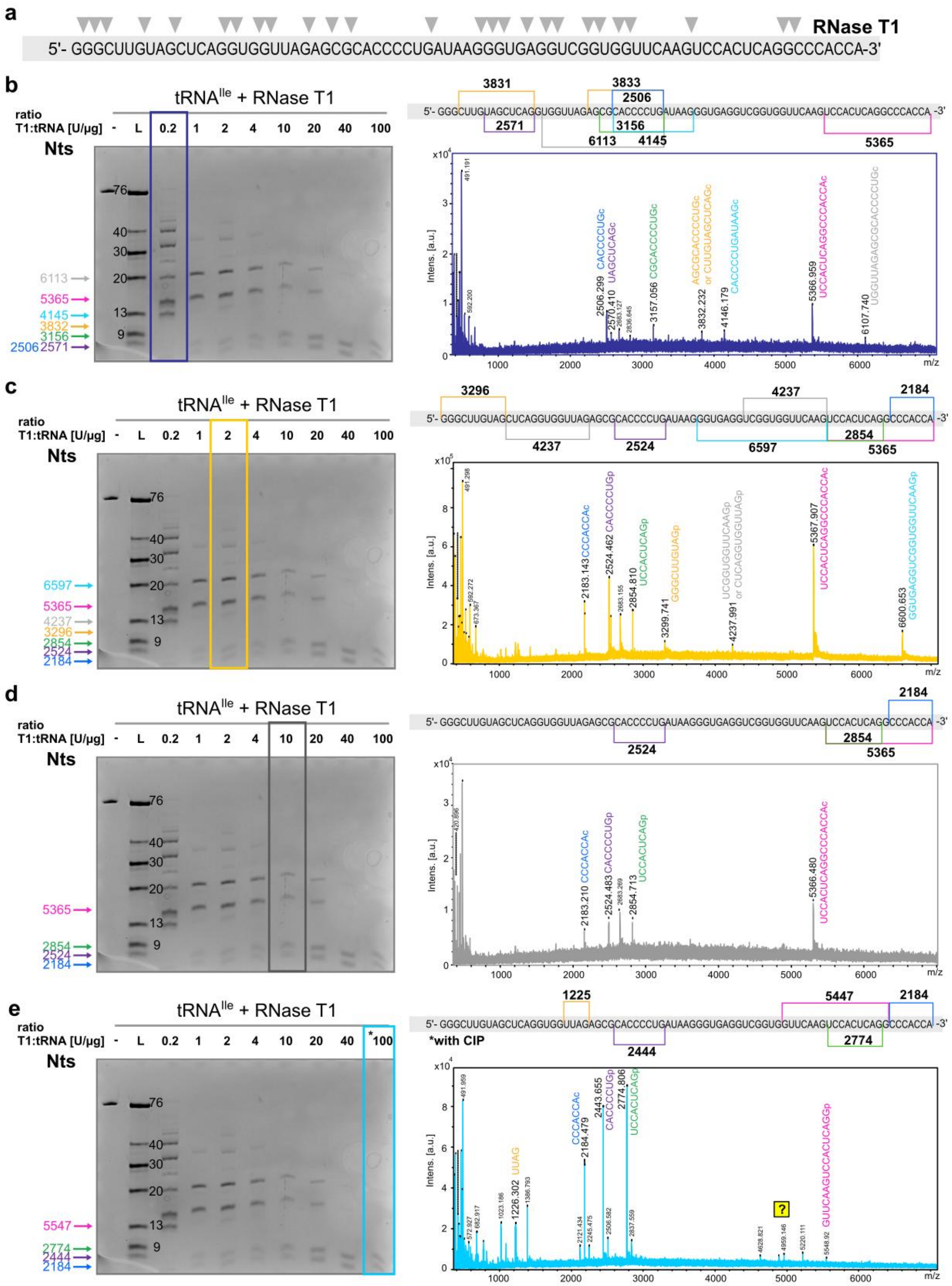

**Figure S2**: Substrate specificity of RNase T1 at different enzyme to substrate ratios. a) sequence of an unmodified tRNA (E. c. tRNA-Ile-GAU) and the expected cleavage sites of RNase T1. b-e) polyacrylamide gel of RNase T1 digested E. c. tRNA-Ile-GAU with different enzyme to substrate ratios and the corresponding MALDI-MS spectra. Assigned fragments and cleavage sites are added to the MS spectrum and in the sequence above.

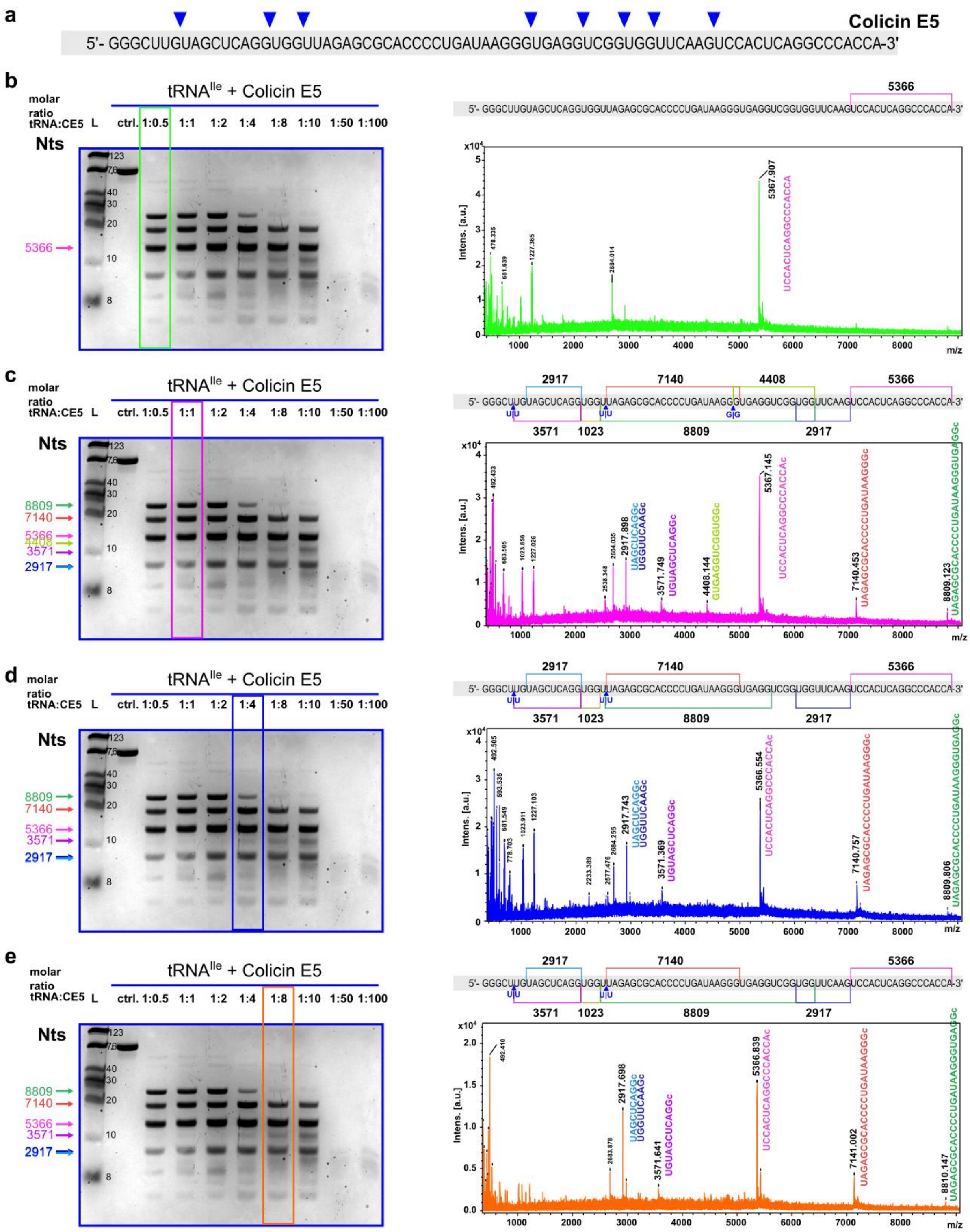

**Figure S3**: Substrate specificity of colicin E5 at different enzyme to substrate ratios. a) sequence of an unmodified tRNA (E. c. tRNA-Ile-GAU) and the expected cleavage sites of colicin E5. b-e) polyacrylamide gel of colicin E5 digested E. c. tRNA-Ile-GAU with different enzyme to substrate ratios and the corresponding MALDI-MS spectra. Assigned fragments and cleavage sites are added to the MS spectrum and in the sequence above.

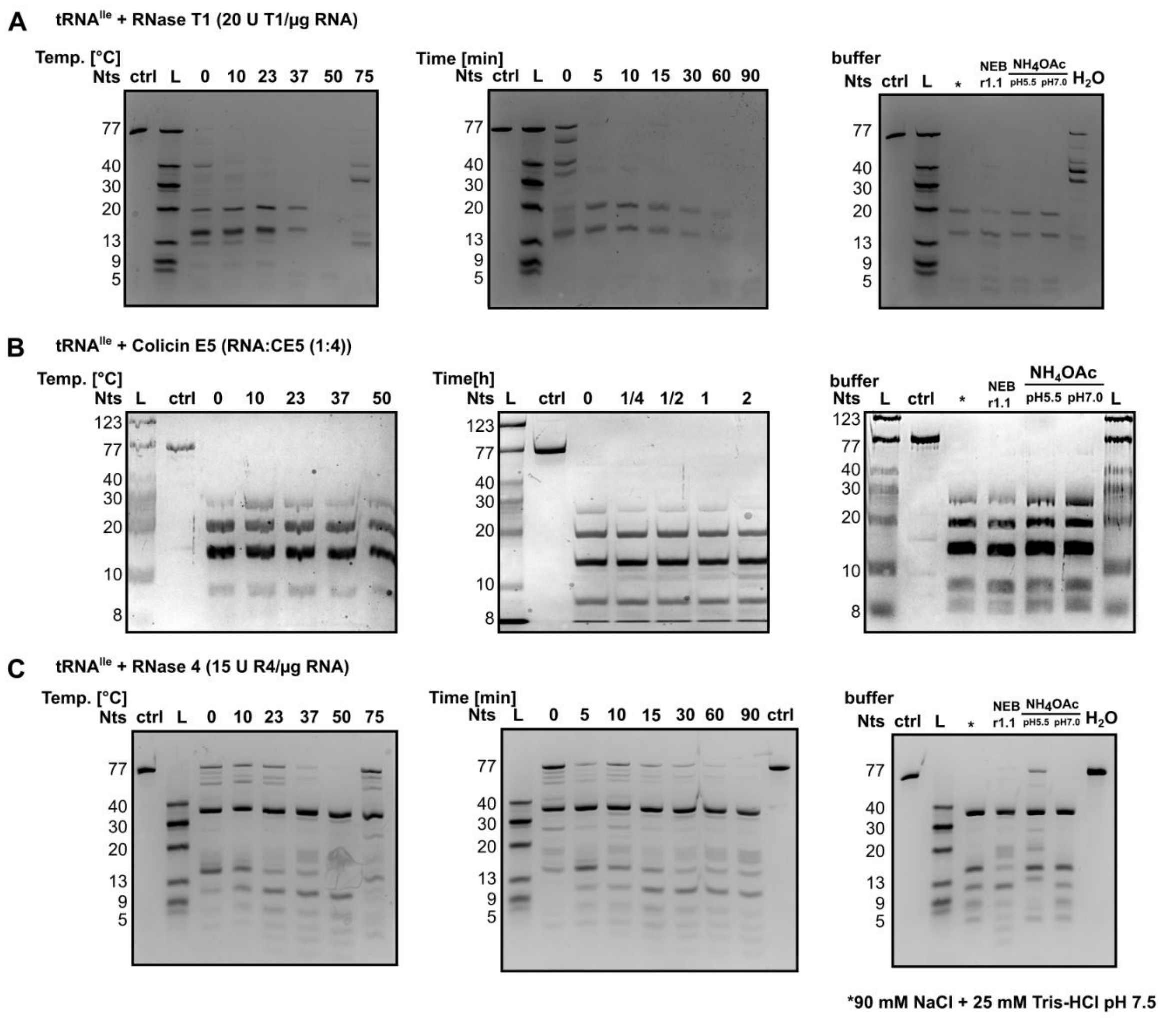

**Figure S4**: Variability of digestion pattern with different reaction conditions. Polyacrylamide gel of E. c. tRNA-Ile-GAU treated with different temperature, incubation time, buffer composition and pH with **A**) RNase T1 **B**) colicin E5 and **C**) RNase 4.

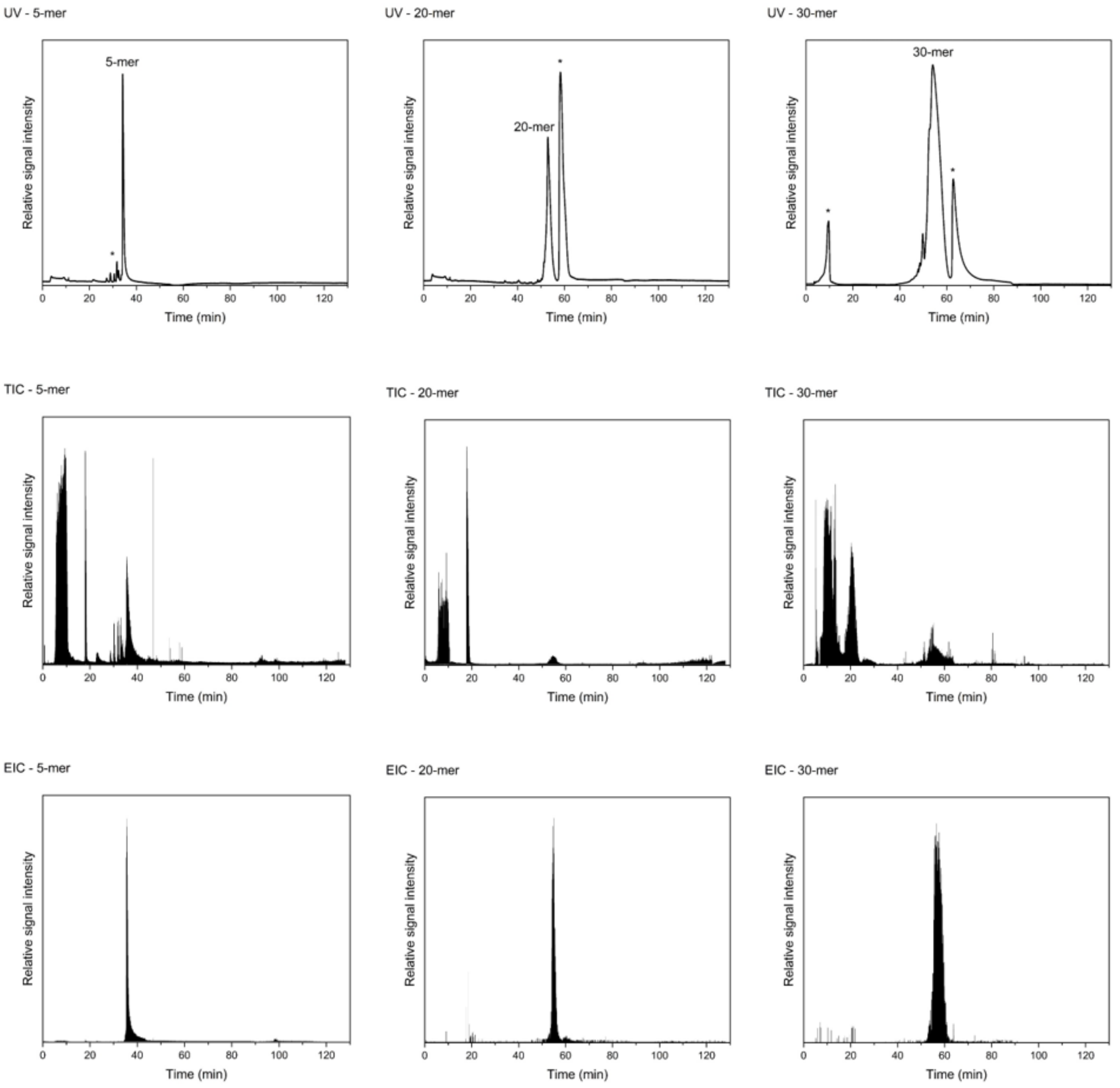

**Figure S5** UV chromatogram, Total ion chromatograms (TICs), and extracted ion chromatograms (EICs) of a 5-mer, a 20-mer, and a 30-mer (synthetic ribonucleotides) analyzed under positive ion mode (POS) via direct injection (DI). Impurities were present in these samples as marked by the asterisk (*) signs in the UV chromatograms.

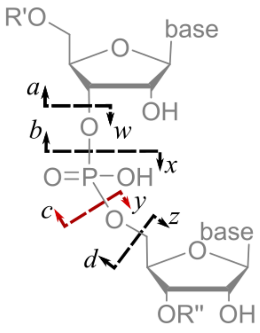

**Figure S6** Nomenclature of MS/MS fragments in oligonucleotide mass spectra. (25)

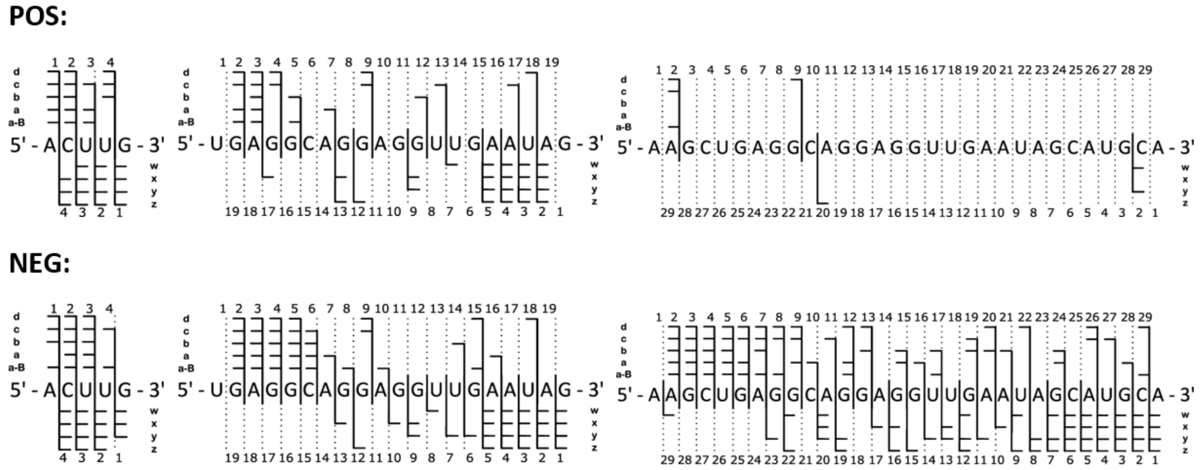

**Figure S7** Sequence coverage of a 5-mer, 20-mer and 30-mer after ionization in positive ion mode (POS) and negative ion mode (NEG). Coverage was determined using NASE (15).

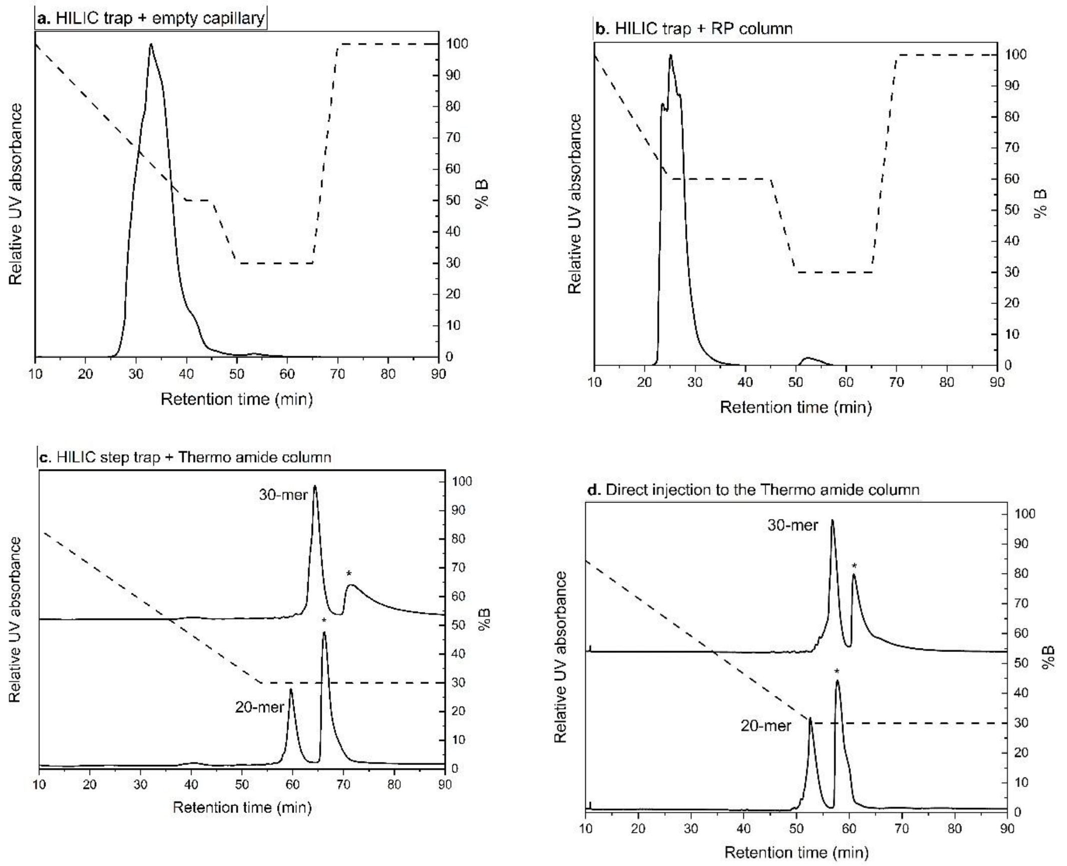

**Figure S8.** Relative UV absorbance at 260 nm of the mixture of a 20-mer and a 30-mer synthetic RNA oligonucleotides (solid line) that had been (5 min at 10 µL/min) trapped on the HILIC stem trap and eluted onto **a.** an empty capillary (150 mm x 0.075 mm), **b.** a nano-flow RP and **c.** a nano-flow HILIC using the "pre-concentration" injection mode. The analysis was repeated using the "direct" injection mode on the **d.** nano-flow HILIC column. The elution flow rates for all analyses were 0.25 µL/min. The corresponding chromatographic gradients (shown as %B) were also indicated as dashed lines. Impurities were present in these samples as marked by the asterisk (*) signs.

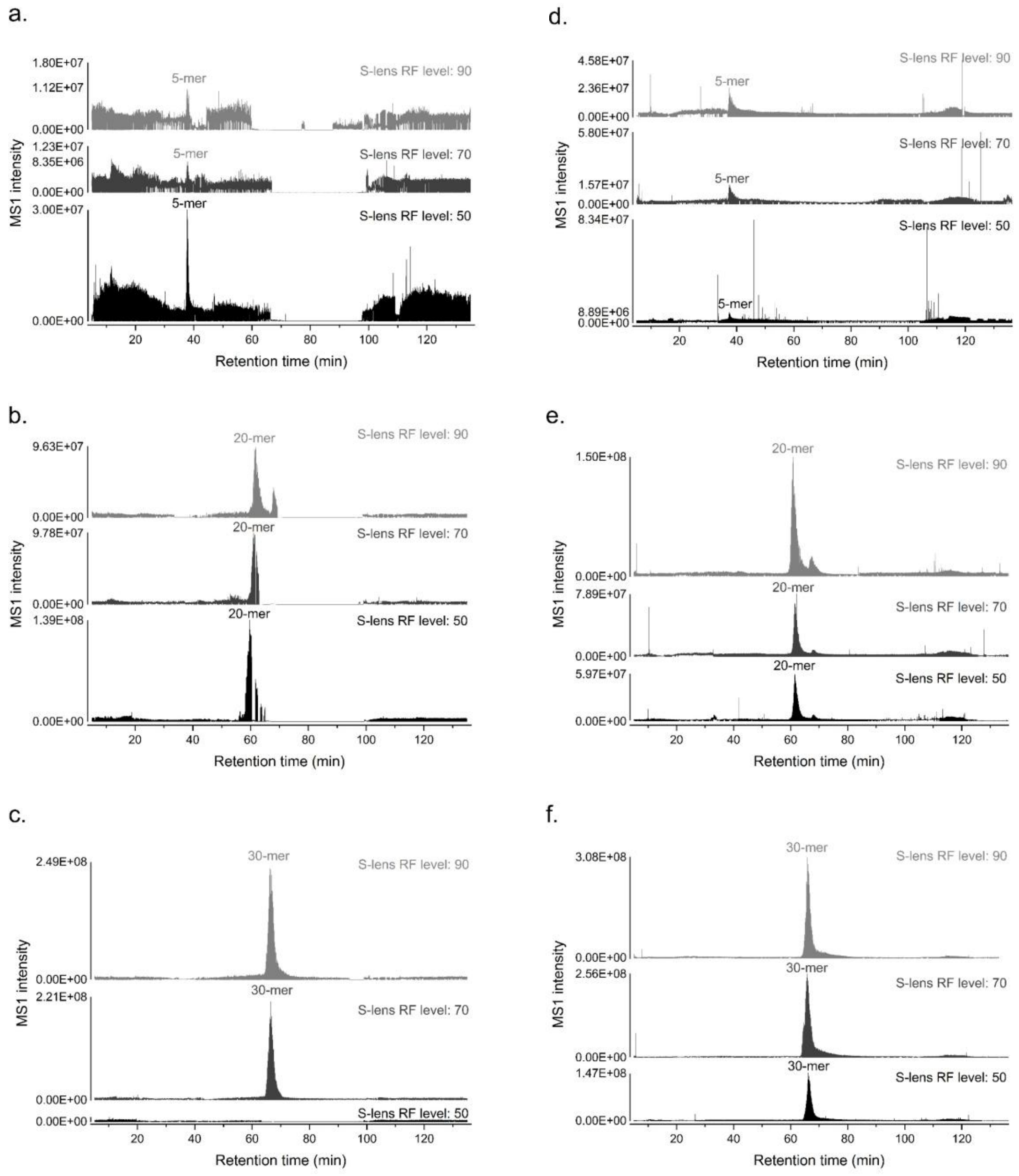

**Figure S9** Total ion chromatograms (TICs) of a 5-mer, 20-mer and a 30-mer (synthetic ribonucleotides) under different S-lens RF levels [i.e. 50 (shown in gray), 70 (shown in dark gray), and 90 (shown in black)] analysed under the regular setup (**a-c**) or the infusion setup (d-e).

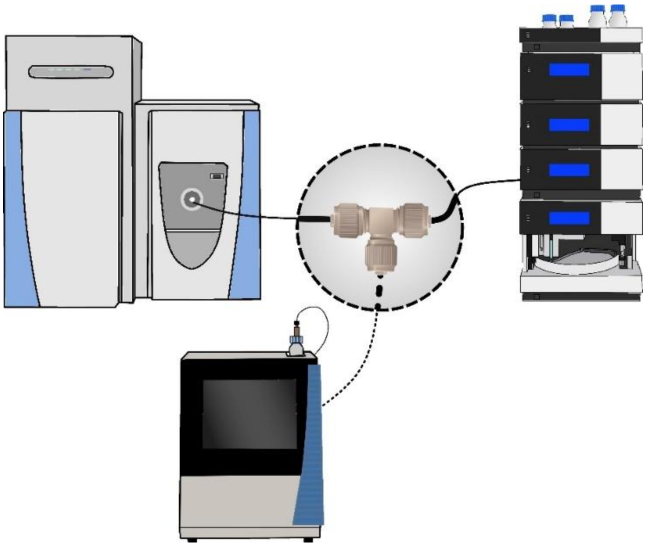

**Figure S10.** Schematic of the instrumental infusion setup. The infusion flow was indicated by the dashed line.

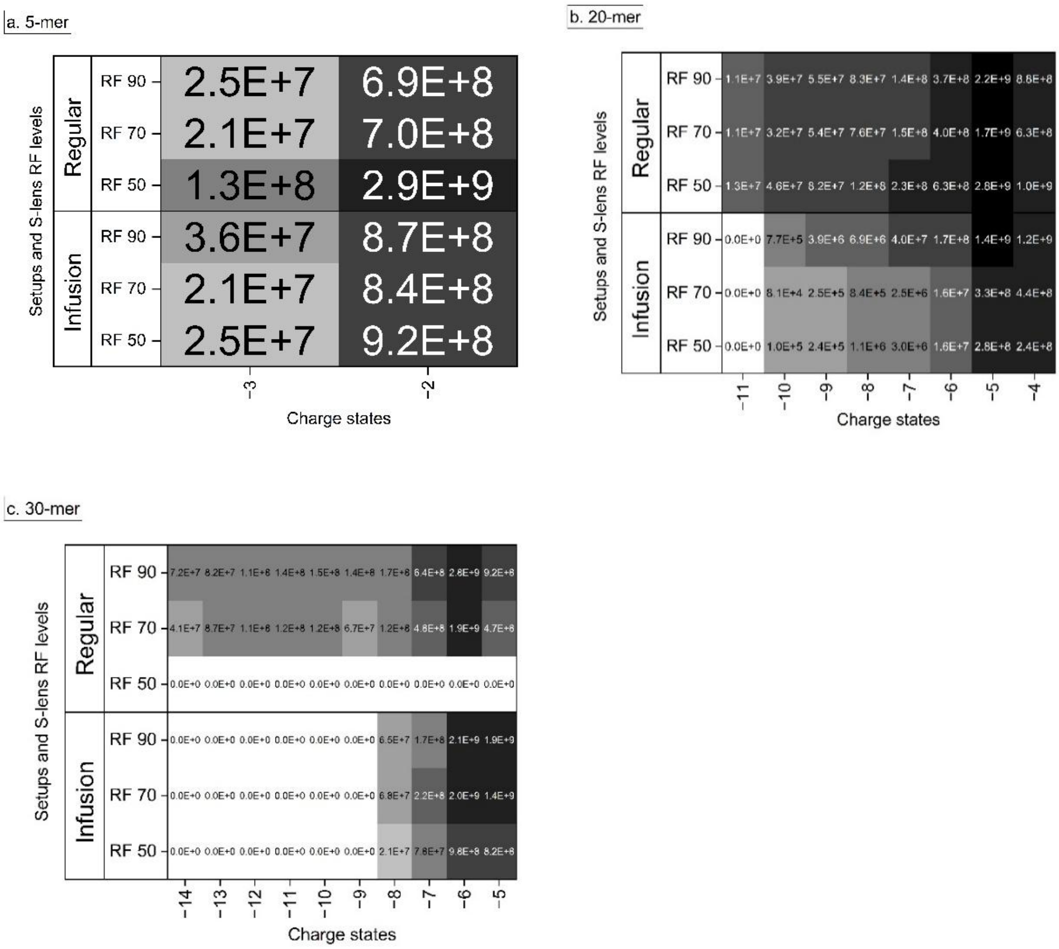

**Figure S11** $MS^1$ signal intensity comparison under different charge states between setups and polarities among **a.** 5-mer, **b.** 20-mer and **c.** 30-mer (synthetic ribonucleotides). The sum of area under curves (AUCs) of the first four isotopic peaks, i.e. [M], [M+1], [M+2], and [M+3], for each charge states were manually integrated in the “skyline” software as the intensity of the corresponding precursor ions.

| **5-mer** | RF 50 | RF 70 | RF 90 |
|---|---|---|---|
| regular | 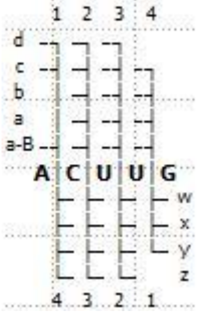 | 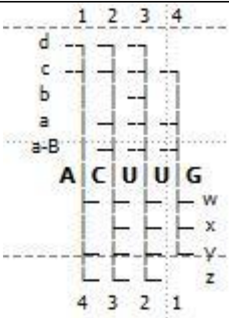 | 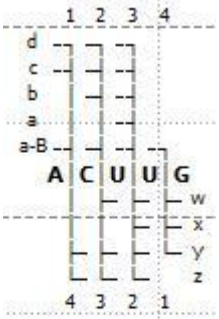 |
| Infusion | 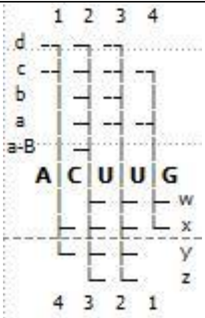 | 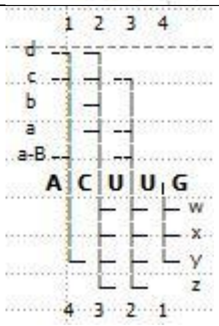 | 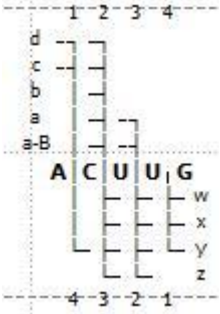 |
| **20-mer** | | | |
| regular | 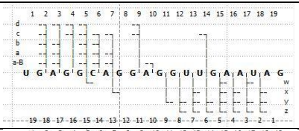 | 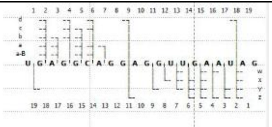 | 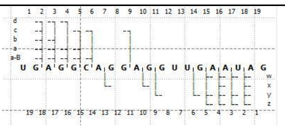 |
| Infusion | 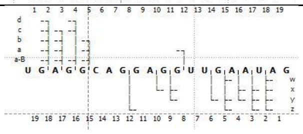 | 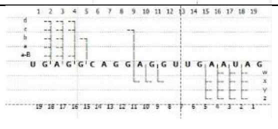 | 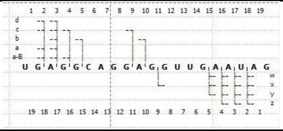 |
| **30-mer** | | | |
| regular | N/A | 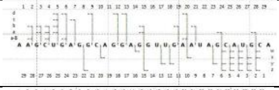 | 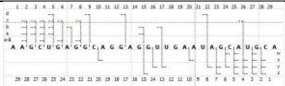 |
| Infusion | 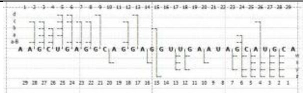 | 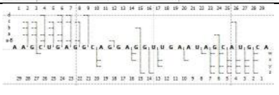 | 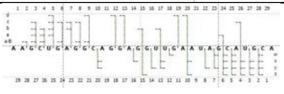 |

**Figure S12.** $MS^2$ sequence coverage comparison between the regular and infusion setup according to NASE analyses. Only the $MS^2$ evidence with the highest matching score was shown as examples.

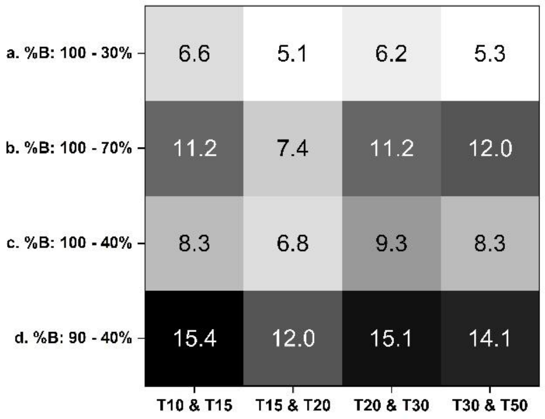

**Figure S13.** Resolutions of indicated peak pairs. **a.** 0-7 min at 100% B, reaches 30% B at 61 min and lasts until 80 min, equilibrates at 100% B from 90 to 140 min. **b.** 0-5 min at 100% B, reaches 40% B at 100 min and lasts until 120 min, equilibrates at 100% B from 130 to 150 min. **c.** 0-5 min at 100% B, reaches 70% B and 40% B at 100 min and 110 min, respectively. After holding at 40% B till 130 min, equilibrates at 100% B from 140 to 170 min. **d.** 0-5 min at 100% B, reaches 90% B, 70% B, and 40% B at 10 min, 100 min and 150 min, respectively. After holding at 40% B till 170 min, equilibrates at 100% B from 180 to 210 min.

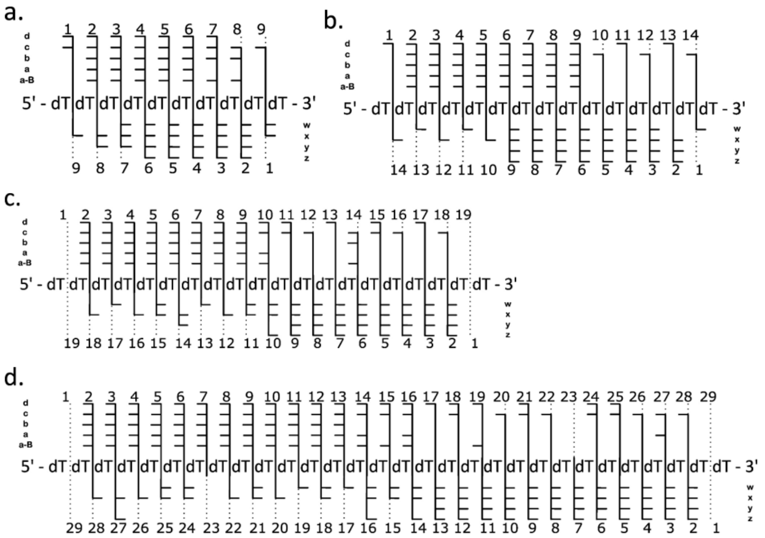

**Figure S14.** MS$^2$ fragmented evidence of **a.** dT10, **b.** dT15, **c.** dT20, and **d.** dT30 identified by NASE among all corresponding MS$^2$ spectra under DDA data acquisition mode from the DNA mixture sample (while the S-lens RF level set at 90). Matched a-B, a, b, c, d, w, x, y, and z ions from the NASE analysis were indicated correspondingly.

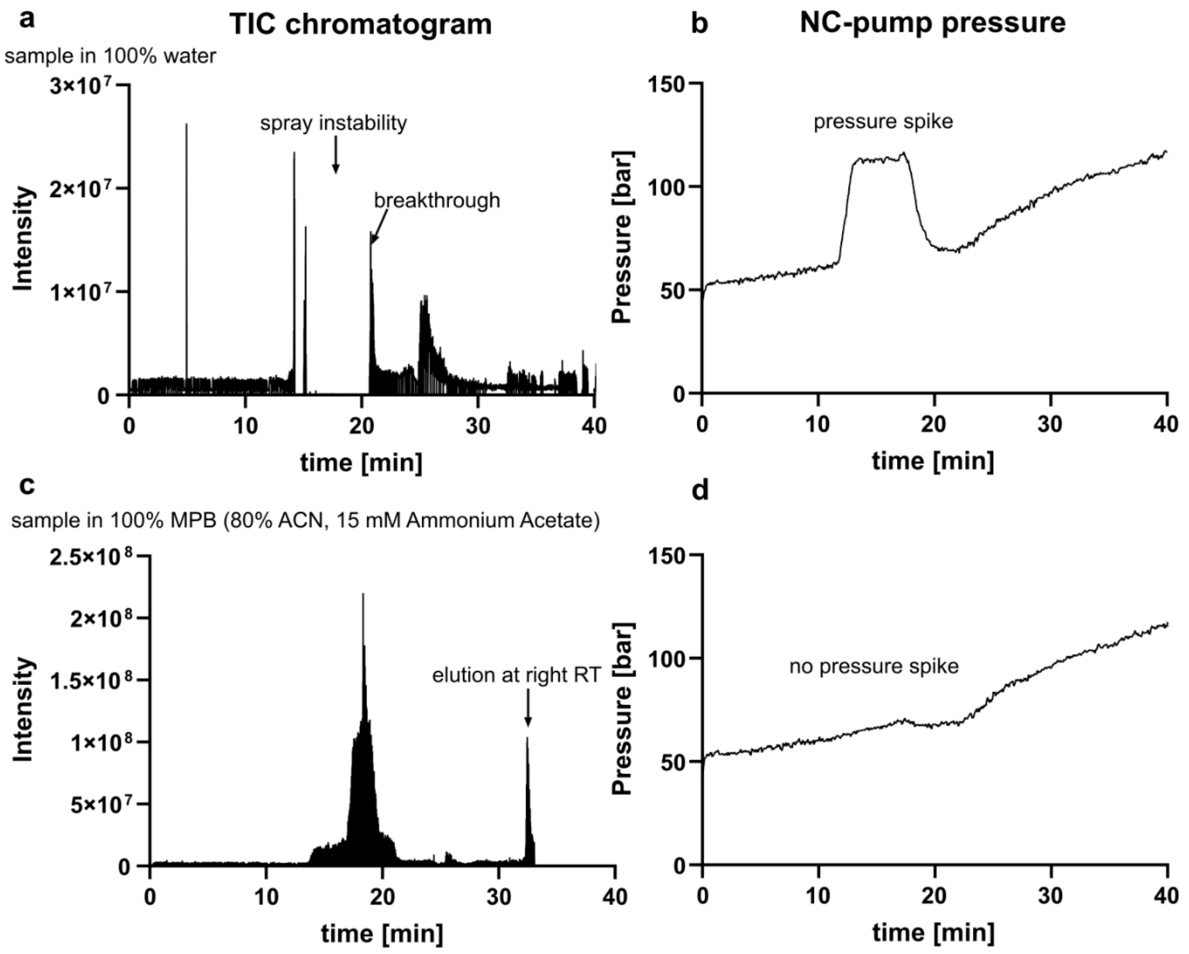

**Figure S15.** TIC and NC-pump trace while direct injection of **a** and **b** 100% aqueous sample, **c** and **d** sample resuspended in 100% mobile phase B (80% ACN, 15 mM ammonium acetate).

tRNA$^{Ile}$ digested by T1 - unspecific cleavage

| Observed m/z | Charge | Precursor error (\|ppm\|) | Theoretical* Molecular mass | Search parameter | Oligonucleotide length | Oligonucleotide sequence | Score | cleavage site | Retention time (min) |
|---|---|---|---|---|---|---|---|---|---|
| 840,11 | -3 | 2,25 | 2396,529 | T1: 3'-P8-mer | | CACCCCUG-P | 275,86 | G\|; G\| | 24,90 |
| 950,46 | -3 | 4,03 | 2710,725 | T1: 3'-P9-mer | | UCCACUCAG-P | 271,85 | G\|; G\| | 25,82 |
| 968,12 | -2 | 2,12 | 1843,170 | T1: 3'-P6-mer | | UUCAAG-P | 161,99 | G\|; G\| | 22,76 |
| 738,43 | -3 | 2,24 | 2107,345 | T1: 3'-P7-mer | | ACCCCUG-P | 138,85 | G\|; G\| | 23,50 |
| 803,10 | -2 | 1,80 | 1528,975 | T1: 3'-P5-mer | | CUCAG-P | 115,30 | G\|; G\| | 23,18 |
| 827,12 | -2 | 2,44 | 1577,025 | T1: 3'-P5-mer | | AUAAG-P | 104,26 | G\|; G\| | 22,90 |
| 732,76 | -3 | 9,38 | 2067,219 | T1: 3'-P7-mer | | CACCCCU-P | 83,47 | G\|; U\| | 22,88 |
| 651,08 | -2 | 9,87 | 1240,775 | T1: 3'-P4-mer | | UUAG-P | 82,68 | G\|; G\| | 21,52 |
| 732,77 | -3 | 9,84 | 2091,345 | T1: 3'-P7-mer | | CCACUCA-P | 80,27 | U\|; A\| | 24,59 |
| 926,12 | -2 | 3,37 | 1801,175 | T1: 3'-P6-mer | | CCACCA-P | 72,04 | C\| | 22,82 |
| 971,12 | -3 | 9,07 | 2710,725 | T1: 3'-P9-mer | | AGUCCACUC-P | 69,67 | A\|; C\| | 23,44 |
| 969,80 | -3 | 0,62 | 2709,740 | T1: 3'-P9-mer | | CACCCCUGA-P | 69,14 | G\|; A\| | 24,08 |
| 968,12 | -2 | 8,90 | 1842,185 | T1: 3'-P6-mer | | ACUCAG-P | 59,79 | C\|; G\| | 24,62 |
| 965,11 | -3 | 6,81 | 2734,750 | T1: 3'-P9-mer | | UCAAGUCCA-P | 58,88 | U\|; A\| | 24,47 |
| 854,76 | -3 | 4,74 | 2438,525 | T1: 3'-P8-mer | | UGUAGCUC-P | 49,34 | U\|; C\| | 24,02 |
| 983,77 | -3 | 9,79 | 2774,775 | T1: 3'-P9-mer | | CCUGAUAAG-P | 47,26 | C\|; G\| | 23,98 |
| 960,46 | -3 | 0,90 | 2725,740 | T1: 3'-P9-mer | | CUCAGGCCC-P | 47,21 | A\|; C\| | 26,29 |
| 841,43 | -3 | 6,46 | 2397,515 | T1: 3'-P8-mer | | CCCCUGAU-P | 42,95 | A\|; U\| | 23,62 |
| 855,10 | -3 | 7,08 | 2420,555 | T1: 3'-P8-mer | | CCACUCAG-P | 40,22 | U\|; G\| | 23,82 |
| 1271,66 | -2 | 3,74 | 2396,529 | T1: 3'-P8-mer | | CACCCCUG-P | 37,25 | G\|; G\| | 24,89 |
| 773,61 | -2 | 1,30 | 1511,990 | T1: 3'-P5-mer | | CACCA-P | 36,71 | C\| | 22,26 |
| 1181,82 | -3 | 7,00 | 3353,145 | T1: 3'-P11-mer | | AGUCCACUCAG-P | 34,52 | A\|; G\| | 22,78 |
| 1286,17 | -2 | 6,13 | 2444,580 | T1: 3'-P8-mer | | CAAGUCCA-P | 34,30 | U\|; A\| | 23,27 |
| 814,09 | -2 | 1,82 | 1528,975 | T1: 3'-P5-mer | | GUCCA-P | 33,33 | A\|; A\| | 23,84 |

tRNA$^{Ile}$ digested by RNase 4 - unspecific cleavage

| Observed m/z | Charge | Precursor error (\|ppm\|) | Theoretical* Molecular mass | Search parameter | Oligonucleotide length | Oligonucleotide sequence | Score | cleavage site | Retention time (min) |
|---|---|---|---|---|---|---|---|---|---|
| 998,64 | -2 | 7,67 | 1842,255 | RNase 4: 2',3'-cP | 6-mer | AAGGGU-cP | 104,53 | U\|A; U\|G | 21,69 |
| 665,42 | -3 | 6,76 | 1842,255 | RNase 4: 2',3'-cP | 6-mer | AAGGGU-cP | 104,21 | U\|A; U\|G | 22,16 |
| 926,62 | -2 | 8,01 | 1698,155 | RNase 4: 2',3'-cP | 6-mer | ACCCCU-cP | 86,43 | C\|A; U\|G | 22,40 |
| 794,10 | -2 | 6,57 | 1448,995 | RNase 4: 2',3'-cP | 5-mer | AGCUC-cP | 100,53 | U\|A; C\|A | 21,74 |
| 661,58 | -2 | 4,89 | 1199,835 | RNase 4: 2',3'-cP | 4-mer | AGGU-cP | 71,84 | C\|A; U\|G | 21,14 |
| 665,09 | -3 | 9,64 | 1779,190 | RNase 4: 2',3'-cP | 6-mer | UCAGGU-cP | 8,40 | C\|U; U\|G | 20,68 |
| 1066,81 | -3 | 0,81 | 2982,010 | RNase 4: 2',3'-cP | 10-mer | AGGCCCACCA-cP | 178,79 | C\|A | 24,62 |

tRNA$^{Ile}$ digested by Colicin E5 - unspecific cleavage

| Observed m/z | Charge | Precursor error (\|ppm\|) | Theoretical* Molecular mass | Search parameter | Oligonucleotide length | Oligonucleotide sequence | Score | cleavage site | Retention time (min) |
|---|---|---|---|---|---|---|---|---|---|
| 971,12 | -3 | 0,92 | 2915,378 | unspecific: 2',3'-cP | 9-mer | UAGCUCAGG-cP | 254,27 | G\|U; G\|U | 23,46 |
| 728,09 | -4 | 1,15 | 2915,378 | unspecific: 2',3'-cP | 9-mer | UAGCUCAGG-cP | 228,61 | G\|U; G\|U | 23,71 |
| 767,10 | -3 | 7,76 | 2304,312 | unspecific: 2',3'-cP | 7-mer | UAGAGCG-cP | 182,39 | U\|U; G\|C | 22,90 |
| 719,42 | -3 | 6,67 | 2161,266 | unspecific: 2',3'-cP | 7-mer | UCCACUC-cP | 174,46 | G\|U; C\|A | 22,83 |
| 944,12 | -3 | 0,74 | 2834,381 | unspecific: 2',3'-cP | 9-mer | CACCCCUGA-cP | 155,03 | G\|C; A\|U | 23,82 |
| 947,12 | -2 | 6,46 | 1896,231 | unspecific: 2',3'-cP | 6-mer | UAGCUC-cP | 126,70 | G\|U; C\|A | 22,27 |
| 868,79 | -3 | 9,58 | 2609,353 | unspecific: 2',3'-cP | 8-mer | AGCUCAGG-cP | 123,69 | U\|A; G\|U | 23,20 |
| 665,42 | -3 | 6,21 | 1999,271 | unspecific: 2',3'-cP | 6-mer | UAAGGG-cP | 123,55 | A\|U; G\|U | 22,37 |
| 950,46 | -3 | 0,20 | 2850,376 | unspecific: 2',3'-cP | 9-mer | CUCAGGCCC-cP | 123,21 | A\|C; C\|A | 24,01 |
| 834,11 | -2 | 5,91 | 1670,218 | unspecific: 2',3'-cP | 5-mer | UGAGG-cP | 118,99 | G\|U; G\|U | 21,92 |
| 806,11 | -2 | 5,14 | 1614,217 | unspecific: 2',3'-cP | 5-mer | UCAAG-cP | 118,85 | U\|U; G\|U | 21,61 |
| 998,64 | -2 | 6,45 | 1999,271 | unspecific: 2',3'-cP | 6-mer | UAAGGG-cP | 97,25 | A\|U; G\|U | 22,22 |
| 959,12 | -2 | 5,74 | 1920,242 | unspecific: 2',3'-cP | 6-mer | UUCAAG-cP | 83,00 | G\|U; G\|U | 21,90 |
| 966,63 | -2 | 7,76 | 1935,253 | unspecific: 2',3'-cP | 6-mer | CUCAGG-cP | 72,88 | G\|C; G\|U | 22,40 |
| 794,10 | -2 | 5,03 | 1590,206 | unspecific: 2',3'-cP | 5-mer | AGCUC-cP | 68,92 | U\|A; C\|A | 21,96 |
| 650,08 | -2 | 9,61 | 1301,160 | unspecific: 2',3'-cP | 4-mer | UCGG-cP | 68,42 | G\|U; G\|U | 20,92 |
| 661,59 | -2 | 8,98 | 1325,171 | unspecific: 2',3'-cP | 4-mer | UGAG-cP | 56,07 | G\|U; G\|G | 20,82 |
| 834,11 | -3 | 3,41 | 2505,329 | unspecific: 2',3'-cP | 8-mer | CACCCCUG-cP | 55,32 | G\|C; G\|A | 23,72 |
| 950,46 | -3 | 3,45 | 2853,375 | unspecific: 3'-P | 9-mer | UCCACUCAG-P | 237,08 | G\|U; G\|G | 24,01 |
| 840,11 | -3 | 3,12 | 2523,339 | unspecific: 3'-P | 8-mer | CACCCCUG-P | 211,36 | G\|C; G\|A | 23,92 |
| 968,13 | -2 | 7,10 | 1938,252 | unspecific: 3'-P | 6-mer | UUCAAG-P | 111,99 | G\|U; G\|U | 22,74 |
| 803,11 | -2 | 7,59 | 1608,216 | unspecific: 3'-P | 5-mer | CUCAG-P | 104,01 | G\|C; G\|G | 22,59 |
| 827,12 | -2 | 6,93 | 1656,239 | unspecific: 3'-P | 5-mer | AUAAG-P | 100,60 | G\|A; G\|G | 22,36 |
| 978,12 | -3 | 6,31 | 2933,388 | unspecific: 3'-P | 9-mer | UAGCUCAGG-P | 69,43 | G\|U; G\|U | 23,29 |
| 971,46 | -3 | 1,12 | 2917,393 | unspecific: 3'-P | 9-mer | CCUGAUAAG-P | 59,23 | C\|C; G\|G | 25,66 |
| 651,08 | -2 | 4,95 | 1304,159 | unspecific: 3'-P | 4-mer | UUAG-P | 58,89 | G\|U; G\|A | 21,60 |
| 950,47 | -3 | 7,84 | 2852,391 | unspecific: 3'-P | 9-mer | CACCCCUGA-P | 56,34 | G\|C; A\|U | 24,53 |
| 863,10 | -3 | 9,26 | 2588,341 | unspecific: 3'-P | 8-mer | UAGCUCAG-P | 50,27 | G\|U; G\|G | 22,99 |

**Figure S16** Comparison of RNAse 4, colicin E5 and Rnase T1 digested unmodified RNA *E. c.* tRNA-Ile-GAU) using updated NASE. Table of found fragments for unspecific digestion for RNase T1, RNase 4 and Colicin E5.

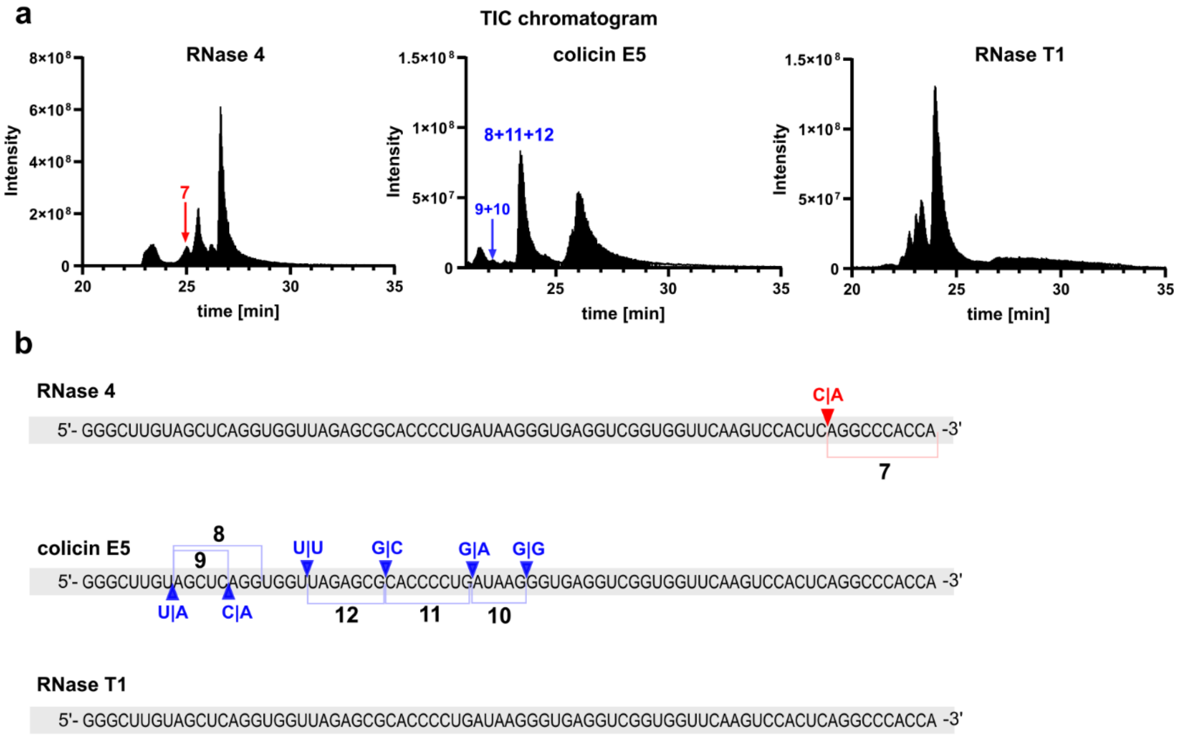

**Figure S17** Unspecific cleavage products found in RNase T1, RNase 4 and colicin E5 digested unmodified RNA (E. c. tRNA-Ile-GAU). **a** TIC with arrows indicating RT of found fragments. **b** Assigned fragments and off-target cleavage sites in the sequence of E. c. tRNA-Ile-GAU. Red: RNase 4; blue: colicin E5; grey: RNase T1.

**tRNA$^{Phe}$ + RNase T1**

| m/z | Observed Charge | Precursor error | Search parameter | Oligonucleotide length | Oligonucleotide sequence | Score | cleavage site | Retention time (min) |
|---|---|---|---|---|---|---|---|---|
| 1400,54 | -3 | 1,55 | T1: 3'-P | 12-mer | A[Cm]U[Gm]AA[yW]AY[m5C]UG-P | 73,28 | G\| | 23,4 |
| 1050,16 | -4 | 4,82 | T1: 3'-P | 12-mer | A[Cm]U[Gm]AA[yW]AY[m5C]UG-P | 62,99 | G\| | 23,45 |

**tRNA$^{Phe}$ + RNase 4**

| m/z | Observed Charge | Precursor error | Search parameter | Oligonucleotide length | Oligonucleotide sequence | Score | cleavage site | Retention time (min) |
|---|---|---|---|---|---|---|---|---|
| 954,50 | -3 | 4,57 | RNase 4: 2',3'-cP | 8-mer | [Gm]AA[yW]AY[m5C]U-cP | 97,18 | U\|Gm; U\|G | 22,84 |
| 983,12 | -3 | 8,70 | RNase 4: 2',3'-cP | 9-mer | A[m2G]CUCAGDD-cP | 76,83 | U\|A; D\|G | 23,11 |
| 1378,66 | -4 | 0,41 | RNase 4: 2',3'-cP | 17-mer | GCGGAUUUA[m2G]CUCAGDD-cP | 12,68 | D\|G | 24,94 |
| 985,81 | -3 | 9,08 | RNase 4: 2',3'-cP | 9-mer | GGAG[m7G]UC[m5C]U-cP | 115,69 | U\|G; U\|G | 24,71 |
| 1338,67 | -4 | 1,87 | RNase 4: 2',3'-cP | 16-mer | GGGAGAGC[m2,2G]CCAGA[Cm]U-cP | 94,34 | D\|G; U\|Gm | 25,30 |
| 959,80 | -3 | 8,92 | unspecific cleavage: 2',3'-cP | 9-mer | CCACAGAAU-cP | 130,95 | U\|A | 23,59 |
| 1189,81 | -3 | 6,66 | unspecific cleavage: 2',3'-cP | 11-mer | G[m5U]YCG[m1A]UCCAC-cP | 80,39 | U\|C; U\|U | 23,97 |

**tRNA$^{Phe}$ + Colicin E5**

| m/z | Observed Charge | Precursor error | Search parameter | Oligonucleotide length | Oligonucleotide sequence | Score | cleavage site | Retention time (min) |
|---|---|---|---|---|---|---|---|---|
| 1454,41 | -5 | 6,67 | Colicin E5: 2',3'-cP | 23-mer | [m5U]YCG[m1A]UCCACAGAAUUCGCACCA | 101,56 | G\|U; G\|U | 25,63 |
| 1297,85 | -2 | 14,00 | Colicin E5: 2',3'-cP | 5-mer | UC[m5C]UG-cP | 96,66 | G\|U; G\|U | 21,63 |
| 754,11 | -3 | 9,38 | Colicin E5: 2',3'-cP | 7-mer | UC[m5C]UGUG-cP | 29,12 | G\|U; G\|U | 22,70 |
| 1055,81 | -3 | 2,77 | unspecific cleavage: 2',3'-cP | 10-mer | [m5U]YCG[m1A]UCCAC-cP | 182,82 | G\|m$^5$U; C\|A | 23,71 |
| 863,78 | -3 | 4,25 | unspecific cleavage: 2',3'-cP | 8-mer | AGAAUUCG-cP | 66,18 | A\|G; G\|C | 23,08 |
| 965,79 | -3 | 1,97 | unspecific cleavage: 2',3'-cP | 9-mer | AGAAUUCGC-cP | 69,12 | A\|G; C\|A | 23,46 |
| 957,78 | -3 | 7,40 | unspecific cleavage: 2',3'-cP | 9-mer | AUUUA[m2G]CUC-cP | 52,89 | G\|A; C\|A | 23,34 |
| 965,79 | -3 | 2,54 | unspecific cleavage: 2',3'-cP | 9-mer | CAGAAUUCG-cP | 49,85 | A\|C; G\|C | 23,64 |
| 1067,14 | -3 | 2,45 | unspecific cleavage: 2',3'-cP | 10-mer | CAGAAUUCGC-cP | 72,68 | A\|C; C\|A | 23,76 |
| 997,81 | -3 | 5,34 | unspecific cleavage: 2',3'-cP | 9-mer | DGGGAGAGC-cP | 131,54 | D\|D; C\|m$^2_2$G | 23,77 |
| 868,11 | -3 | 6,43 | unspecific cleavage: 2',3'-cP | 8-mer | G[m1A]UCCACA-cP | 89,88 | C\|G; A\|G | 23,17 |
| 973,79 | -3 | 7,40 | unspecific cleavage: 2',3'-cP | 9-mer | GAAUUCGCA-cP | 59,73 | A\|G; A\|C | 23,33 |
| 1081,15 | -3 | 7,56 | unspecific cleavage: 2',3'-cP | 10-mer | GAAUUCGCAC-cP | 65,74 | A\|G; C\|C | 23,81 |
| 860,45 | -3 | 2,99 | unspecific cleavage: 2',3'-cP | 8-mer | UA[m2G]CUCAG-cP | 211,38 | U\|U; G\|D | 22,96 |
| 1062,47 | -3 | 1,15 | unspecific cleavage: 2',3'-cP | 10-mer | UCCACAGAAU-cP | 73,30 | m$^1$A\|U; U\|U | 23,57 |
| 1379,21 | -3 | 2,50 | unspecific cleavage: 2',3'-cP | 8-mer | UCGCACCA | 194,52 | U\|U | 23,18 |

**Figure S18** Comparison of RNase 4, colicin E5 and RNase T1 digested native RNA *S. c.* tRNA-Phe-G$_m$AA) using updated NASE. Table of found fragments for RNase T1, RNase 4 and Colicin E5.

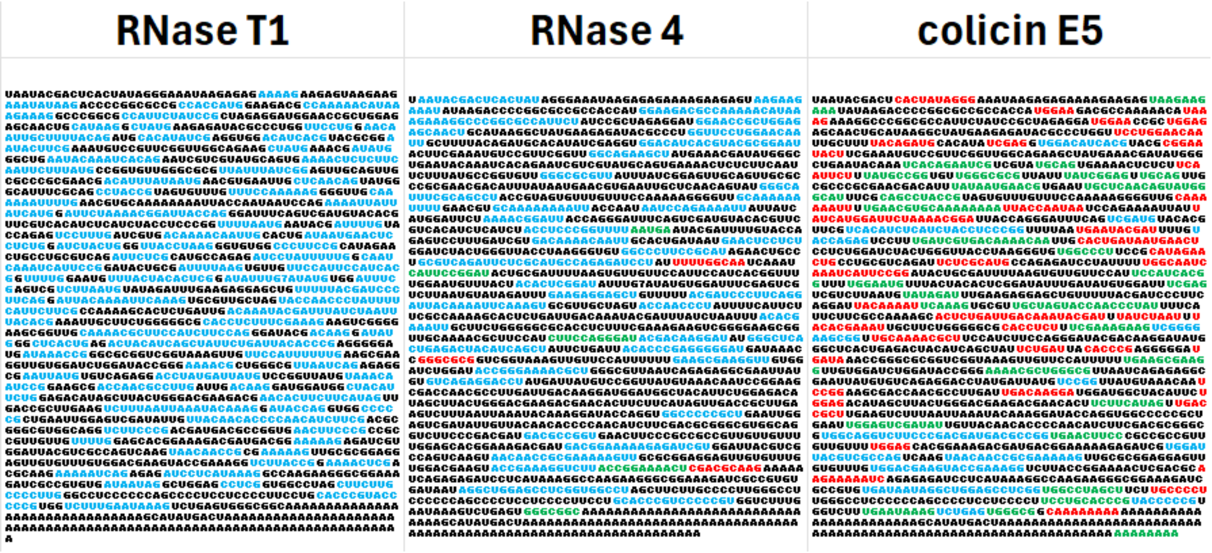

**Figure S19** Comparison of RNase 4, colicin E5 and RNase T1 digested FLuc-mRNA from NIST using updated NASE. Blue fragments: Expected cleavage site on both ends, green fragments: One expected and one unexpected cleavage site and red fragments: Both cleavage sites are unexpected.

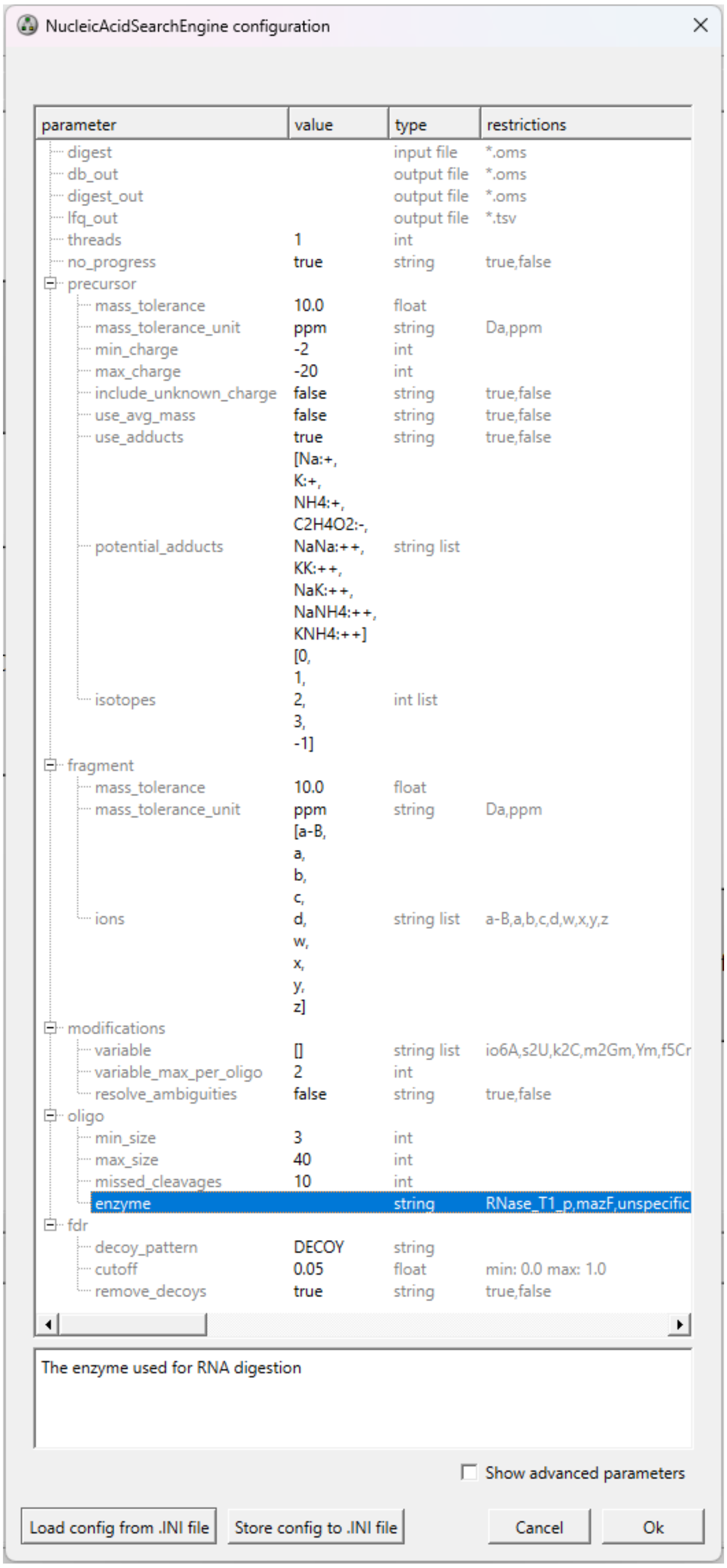

**Figure S20** NASE configuration for the data analysis of IVT and biological RNA samples. The right nuclease is chosen for the row “enzyme”, which is highlighted in blue.
